# Preparation of *E. coli* RNA polymerase transcription elongation complexes for systematic RNA assays

**DOI:** 10.1101/2021.03.15.435517

**Authors:** Eric J. Strobel

## Abstract

RNA folds into secondary and tertiary structures that can mediate diverse cellular functions. Understanding how RNA sequence directs the formation of biologically active structures requires approaches that can comprehensively assess how changes in an RNA sequence affect its structure and function. Towards this goal, I have developed a general method for purifying *E. coli* RNA polymerase (RNAP) transcription elongation complexes (TECs) for use in systematic RNA assays. My approach depends on two constituent technologies: First, I have designed an *E. coli* σ ^70^ promoter that can be efficiently barcoded using a one-pot series of enzymatic reactions. Second, I have developed a strategy for purifying promoter-initiated *E. coli* RNAP TECs by selective photo-elution from streptavidin-coated magnetic beads. Together, these methods establish a platform for the development of TEC Display assays in which the functional properties of RNA sequence variants can be recorded by fractionating and quantitatively barcoding a TEC library.

## Introduction

RNA molecules perform diverse cellular functions that frequently depend on secondary and tertiary structures (1). RNA structure formation is dynamic, complex, and begins when RNA is synthesized during transcription (2-4). Understanding how RNA primary sequence gives rise to biologically active structures therefore requires quantitative methods that can efficiently explore the functional capacity of vast sequence space, cotranscriptionally.

One strategy for comprehensively assessing the function of complex RNA sequence libraries is to couple high-throughput DNA sequencing with *in situ* transcription on an Illumina sequencer flow cell to perform fluorescence-based RNA assays (5). This approach, which was pioneered by the <u>Hi</u>gh-<u>T</u>hroughput <u>S</u>equencing-<u>R</u>NA <u>A</u>ffinity <u>P</u>rofiling (HiTS-RAP) (6) and Quantitative Analysis of <u>RNA</u> on a <u>Ma</u>ssively <u>P</u>arallel Array (7) methods and has collectively been referred to as <u>RNA</u> array on a <u>Hi</u>gh-<u>T</u>hroughput <u>S</u>equencer (RNA-HiTS) (5), leverages the ability of stalled TECs to physically link an RNA transcript to the DNA template from which it was transcribed. In this way, direct measurements of RNA function can be coupled to a DNA sequencing read.

Independent of its technical utility for linking transcribed RNA to its DNA template in RNA-HiTS assays, the cotranscriptional display of RNA is a powerful tool for studies of RNA folding for several reasons. First, RNA molecules begin to fold during transcription (3,4) and cotranscriptionally folded RNA may adopt different structures than renatured RNA (8-10). Second, experiments in which RNA is tethered to its DNA template can circumvent biases from processing steps that are typically required for RNA sequencing libraries, such as ssRNA ligation and reverse transcription, because sequence information can be recovered from the template DNA. Similarly, the presence of a physical link between RNA function and DNA sequence can enable the investigation of intrinsically destructive RNA activities, such as self-cleavage (11), in which RNA sequence information is lost when the RNA performs its biochemical function. Third, direct interactions between a transcribing RNAP and nascent RNA are frequently required for cotranscriptional RNA functions (12-14). Cotranscriptional RNA assays are therefore essential to our understanding of RNA structure and function.

The application of RNA-MaP and HiTS-RAP to diverse RNA targets illustrates the utility of cotranscriptionally displayed RNA libraries for uncovering the sequence and structural basis of RNA function (6,7,11,15-20). Nonetheless, TEC-based RNA array experiments in which the measurement of RNA function is independent of a sequencing instrument have not been developed. In contrast to RNA-HiTS, an off-sequencer TEC-based RNA assay would necessarily sacrifice the ability to directly measure RNA function. However, an appropriately designed method would retain many of the advantages gained by coupling RNA to DNA, would be accessible to laboratories without the technical expertise for RNA-HiTS assays, and could likely be integrated with other biochemical assays such as RNA structure probing (21) and *in vitro* selection (22,23). An off-sequencer platform for the development of TEC-based RNA assays would therefore be distinct from and complementary to the established RNA-HiTS methodologies.

In this work, I have developed a <u>T</u>ranscription <u>E</u>longation <u>C</u>omplex Display (TEC Display) method that combines TEC purification with quantitative DNA barcoding to establish a platform for the development of systematic RNA assays. First, I describe a derivative of the λP_R_ σ^70^ promoter that can be efficiently barcoded in a one-pot, sequence-independent series of enzymatic reactions. I then implement a TEC purification procedure based on my previous observation that stalling *E. coli* RNAP at an internal desthiobiotin-TEG (or biotin-TEG, as described here) lesion blocks attachment of the biotinylated DNA to streptavidin-coated magnetic beads (24). In this procedure, a DNA template containing both an internal biotin-TEG stall site and a 5’ <u>p</u>hoto<u>c</u>leavable biotin (5’ PC biotin) is transcribed and attached to streptavidin beads. DNA containing TECs is attached by PC biotin only and can therefore be selectively photo-eluted, whereas DNA without a TEC is retained by the exposed biotin-TEG stall site. My procedures for TEC purification and quantitative DNA barcoding can be integrated seamlessly, and together will enable the development of experiments in which measurements of RNA function are directly coupled to DNA sequence using a TEC library.

## Results

### Rationale and requirements for TEC Display experiments

I developed the *E. coli* σ^70^ promoter barcoding and TEC purification methods presented in this work to facilitate *in vitro* transcription assays in which a DNA library can be fractionated by the function of cotranscriptionally displayed RNA. In principle, barcoding the partitioned DNA library with fraction- and molecule-specific sequence identifiers can quantitatively record the activity of each RNA variant as the distribution of its corresponding template DNA between fractions. A successful implementation of this strategy requires that the barcoding reaction is 1) DNA sequence-independent so that it is compatible with complex DNA libraries, 2) efficient and minimally biased, and 3) coupled with a procedure to purify TECs so that only DNA templates that have directed productive transcription are used in the assay. The procedure I have developed achieves these criteria using an extensively modified linear DNA template that contains the following features: First, an internal biotin-TEG modification functions as a chemically encoded stall site for *E. coli* RNAP (24) and as an attachment point, first for depleting DNA without TECs and subsequently for purifying barcoded DNA; the internal biotin-TEG is embedded in a variant of the Illumina TruSeq Small RNA RP1 primer that was extended by five nucleotides so that it contains the full Illumina RA5 adapter sequence. Second, thermostability-enhancing dNTPs ensure DNA integrity downstream of the internal biotin-TEG modification during DNA barcoding. Third, a 5’ PC biotin enables reversible DNA immobilization for buffer exchanges during TEC purification. Fourth, two deoxyuridine (dU) nucleotides in the promoter can be excised to generate a long 3’ overhang for molecular barcoding. The complete layout of a TEC Display DNA template is shown in Figure 1 and the function of sequence elements and DNA modifications are described in detail in the relevant sections below.

**Figure 1.**
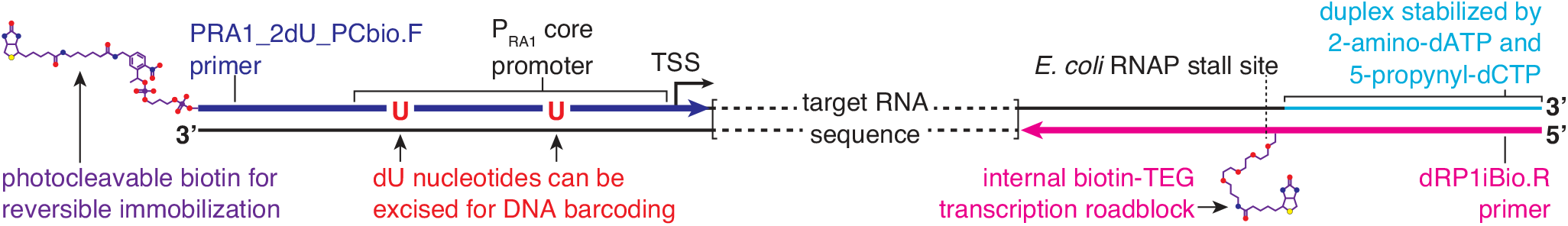
Layout of chemically modified constant regions used in TEC Display. TEC Display uses two chemically modified constant DNA sequences for integrated TEC purification and DNA barcoding procedures. An internal biotin-TEG modification, which functions as an *E. coli* RNAP stall site and an attachment point during TEC purification and DNA barcoding, is introduced by the dRP1iBio.R primer. Thermostability-enhancing 2-amino-dATP and 5-propynyl-dCTP nucleotides are added to the non-transcribed DNA strand downstream of the internal biotin-TEG modification during the translesion synthesis step of DNA template preparation to stabilize this 29 bp region. The PRA1_2dU_PCbio.F primer introduces a 5’ PC biotin modification for reversible DNA immobilization during TEC purification along with two dU nucleotides that are excisable to facilitate DNA barcoding. Oligonucleotide sequences, including information about required purifications, are available in Table S1.

### dU excision enables a one-pot dsDNA barcoding reaction

dU excision is an established method for generating ‘sticky’ ends in molecular cloning protocols (25). In this approach, the thermolabile <u>U</u>racil-<u>S</u>pecific <u>E</u>xcision <u>R</u>eagent (USER^®^) enzyme excises dU nucleotides from DNA through the activity of uracil DNA glycosylase (26,27), which catalyzes excision of the uracil base, and endonuclease III which excises the resulting abasic site leaving a one nucleotide gap with a 5’ phosphate (28,29). To facilitate USER^®^-mediated promoter barcoding, I designed the P_RA1_ promoter which is derived from the bacteriophage promoters λ<u>P_R_</u> and T7<u>A1</u> (Fig. 2A). I selected λP_R_ as the primary basis (positions -1 to -35) of P_RA1_ due to the long lifetime of λP_R_ open promoter complexes (30). Sequence upstream of -35 (−36 to -50) comprises the proximal UP element subsite (31) of the T7A1 promoter to reduce the GC content of this region. I positioned dU nucleotides at -13 (one nucleotide upstream of the -10 element hexamer) and at -30 (the sixth nucleotide in the -35 hexamer). These positions were selected purposefully so that to the Tm of the DNA fragments that are formed upon USER^®^ digestion is dramatically lower than that of resulting 3’ overhang in the transcribed DNA strand (Fig. 2A). -29A was changed to C to promote efficient USER^®^ digestion (32).

**Figure 2.**
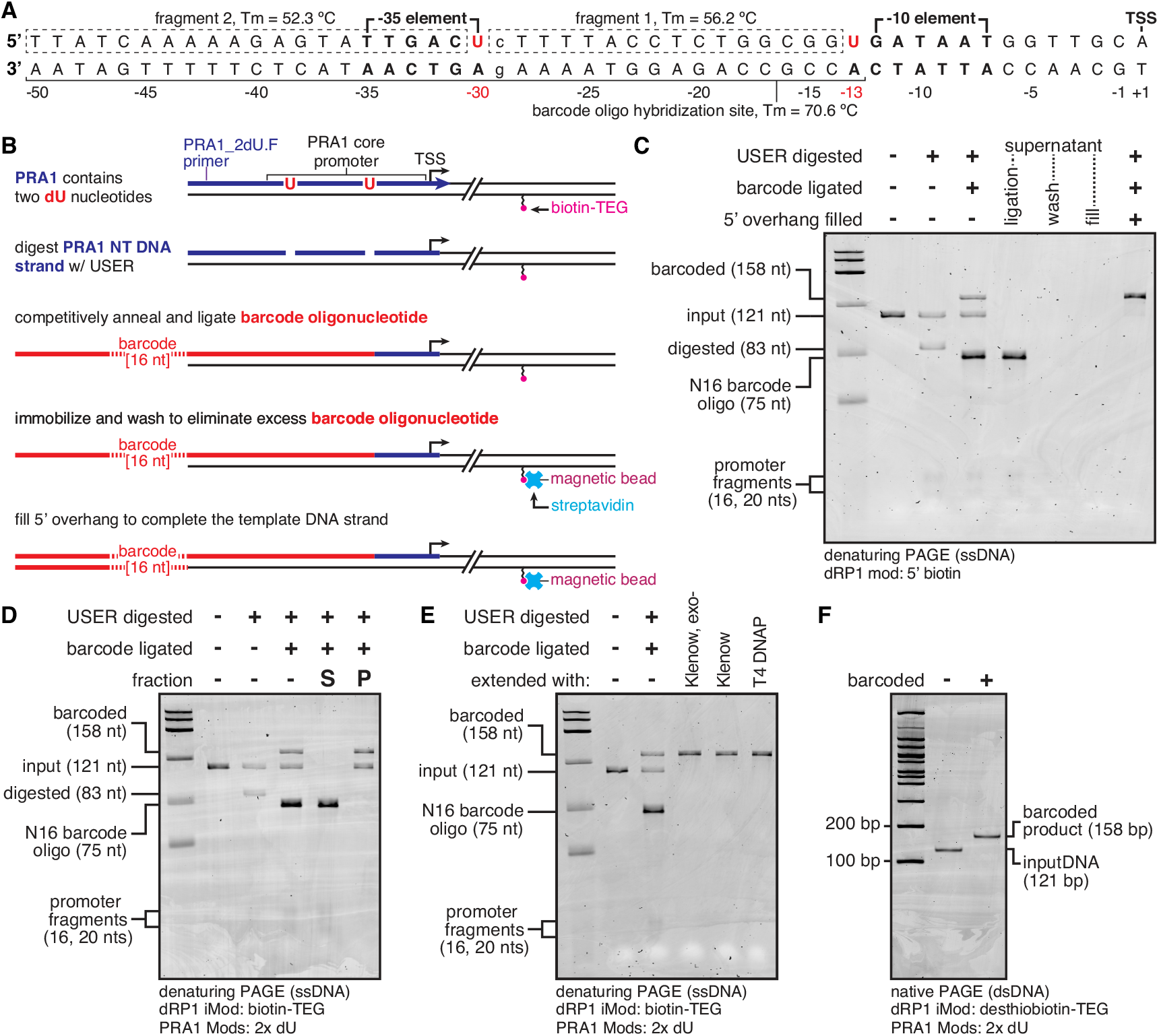
One-pot σ _70_ promoter barcoding. **(A)** Overview of the P_RA1_ promoter. Tm for each fragment and the barcode oligo hybridization site were calculated using the Integrated DNA Technologies OligoAnalyzer Tool with the settings: Oligo conc. = 0.05 μM, Na^+^ conc.= 50 mM, Mg^++^ conc. = 10 mM, dNTPs conc. = 0 mM. **(B)** Overview of the P_RA1_ barcoding procedure. dU nucleotides are excised so that a barcoding oligo can be annealed and ligated to the DNA template. The DNA template is immobilized on streptavidin beads so that excess barcoding oligo can be removed before primer extension to fill the 5’ overhang. **(C)** Proof-of-principle for P_RA1_ barcoding using a 5’ biotinylated DNA. **(D)** Validation of P_RA1_ barcoding with an internal biotin-TEG-modified DNA template through the DNA immobilization step. **(E)** Comparison of DNA polymerases for filling a 5’ overhang on an internal biotin-TEG-modified DNA template. **(F)** Native PAGE comparing input DNA to barcoded DNA. All gels are representative of two independent replicates.

The P_RA1_ promoter barcoding procedure is as follows (Fig. 2B): First, dU nucleotides in the non-transcribed DNA strand of P_RA1_ are excised by digestion with Thermolabile USER^®^ II enzyme. Following digestion, a barcoding oligonucleotide is competitively annealed to the P_RA1_ transcribed DNA strand by heating to 70 °C and slowly cooling to 25 °C. The barcoding oligonucleotide used for protocol validation contains an N16 ‘barcoding region’ flanked by P_RA1_ sequence spanning -50 to -13 at the 3’ end, and the reverse complement of the Illumina TruSeq Small RNA RA3 adapter at the 5’ end (Table S1, PRA1m12_VRA5_16N). Thermolabile USER^®^ II is inactivated during this annealing step. Next, T4 DNA ligase is added to the reaction to ligate the barcoding oligonucleotide to the DNA template and is subsequently heat inactivated. After ligation, the one-pot DNA barcoding procedure is complete and, in experiments that contain fractionated samples, fractions can be pooled so that all downstream manipulations are identical. Excess barcoding oligonucleotide is removed by immobilizing the biotinylated template DNA on streptavidin-coated magnetic beads, removing the supernatant and washing once. The DNA template is then extended to fill the 5’ overhang generated by the barcoding oligonucleotide. Figure 2C shows a proof-of-principle barcoding reaction using a 5’ biotinylated DNA template (DNA Template 1 in Table S2), in which one sample volume was removed after key manipulations to visualize step-by-step processing. Each step is efficient, and no reaction side products are detectable.

### P_RA1_ barcoding is compatible with internally modified DNA

I previously described a method for preparing internally modified linear dsDNA that can be used to halt transcription elongation by *E. coli* RNAP for cotranscriptional RNA assays (24). This *chemical transcription roadblocking* approach enables an RNAP stall site to be functionalized for complex manipulations of the stalled TECs. My protocol for the sequence-independent preparation of internally modified DNA templates uses the commercially available translesion DNA polymerase *Sulfolobus* DNA polymerase IV (33), which adds 1-2 dA nucleotides to the DNA 3’ ends (34). In principle, a 3’-dA overhang could interfere with primer extension during P_RA1_ barcoding. I therefore assessed the compatibility of P_RA1_ barcoding with a DNA template that contains an internal biotin-TEG modification (DNA Template 2 in Table S2). After demonstrating that internally biotinylated DNA is efficiently barcoded through the oligonucleotide ligation step (Fig. 2D), I performed the final primer extension step using Klenow Fragment (3’→5’ exo-), Klenow Fragment, and T4 DNA polymerase (Fig. 2E). Because primer extension was efficient for each DNA polymerase, I used Klenow Fragment for all subsequent barcoding experiments due to its proofreading capability and because it is less prone to the formation of 3’ recessed ends than T4 DNA polymerase.

Another consideration when barcoding internally modified DNA is whether the relatively short (29 bp) DNA duplex downstream of the modification site remains stable throughout processing, particularly during steps with elevated temperatures such as oligonucleotide annealing. My standard protocol for the preparation of internally modified DNA now uses the thermostability enhancing dNTPs 2-amino-dATP and 5-propynyl-dCTP during translesion synthesis as a safeguard against DNA end-fraying downstream of the modification site. To assess whether undesirable DNA species form during P_RA1_ barcoding, I processed an internally desthiobiotinylated DNA template (DNA Template 3 in Table S2) so that immobilized DNA could be gently eluted from streptavidin-coated magnetic particles by adding free biotin. I observed a complete shift of the dsDNA to the expected length after barcoding and did not observe other DNA species (Fig. 2F), indicating that internally modified dsDNA is a compatible substrate for the P_RA1_ barcoding procedure.

### Purification of precisely positioned TECs by selective photo-elution from streptavidin beads

Transcription initiation is not 100% efficient (35-37). Consequently, TEC Display experiments must be designed such that DNA templates that do not direct productive transcription are excluded from the assay. To this end, I developed a procedure for purifying *E. coli* RNAP TECs based on my previous observation that positioning RNAP at an internal desthiobiotin-TEG stall site blocks attachment to streptavidin-coated magnetic beads (24). To verify that positioning a TEC at an internal *biotin*-TEG stall site blocks streptavidin binding, I assessed the efficiency at which DNA containing an internal biotin-TEG modification was attached to streptavidin beads without and with a single round of transcription. When NTPs were omitted from the transcription reaction, ∼97% of DNA was present in the bead pellet regardless of RNAP concentration, indicating that non-specific binding by excess RNAP did not interfere with DNA immobilization (Fig. 3A, B). The ∼3% of DNA that is not attached to the streptavidin beads likely reflects a DNA population that does not contain the biotin modification or in which the biotin modification was inactivated during oligonucleotide synthesis. When NTPs were included to permit single-round transcription, ∼55% of DNA was excluded from the streptavidin beads in all conditions where RNAP was included (Fig. 3B). Consistent with the interpretation that streptavidin bead exclusion is TEC-dependent, the 42 nt RNA transcript that is produced when RNAP is halted at the internal biotin stall site was only observed in the supernatant (Fig. 3B). Given that the DNA template was saturated by open complexes and that transcription is single-round, the exclusion of ∼52% of DNA above background suggests that, under the conditions of this assay, approximately half of P_RA1_ promoter open complexes proceed to productive elongation.

**Figure 3.**
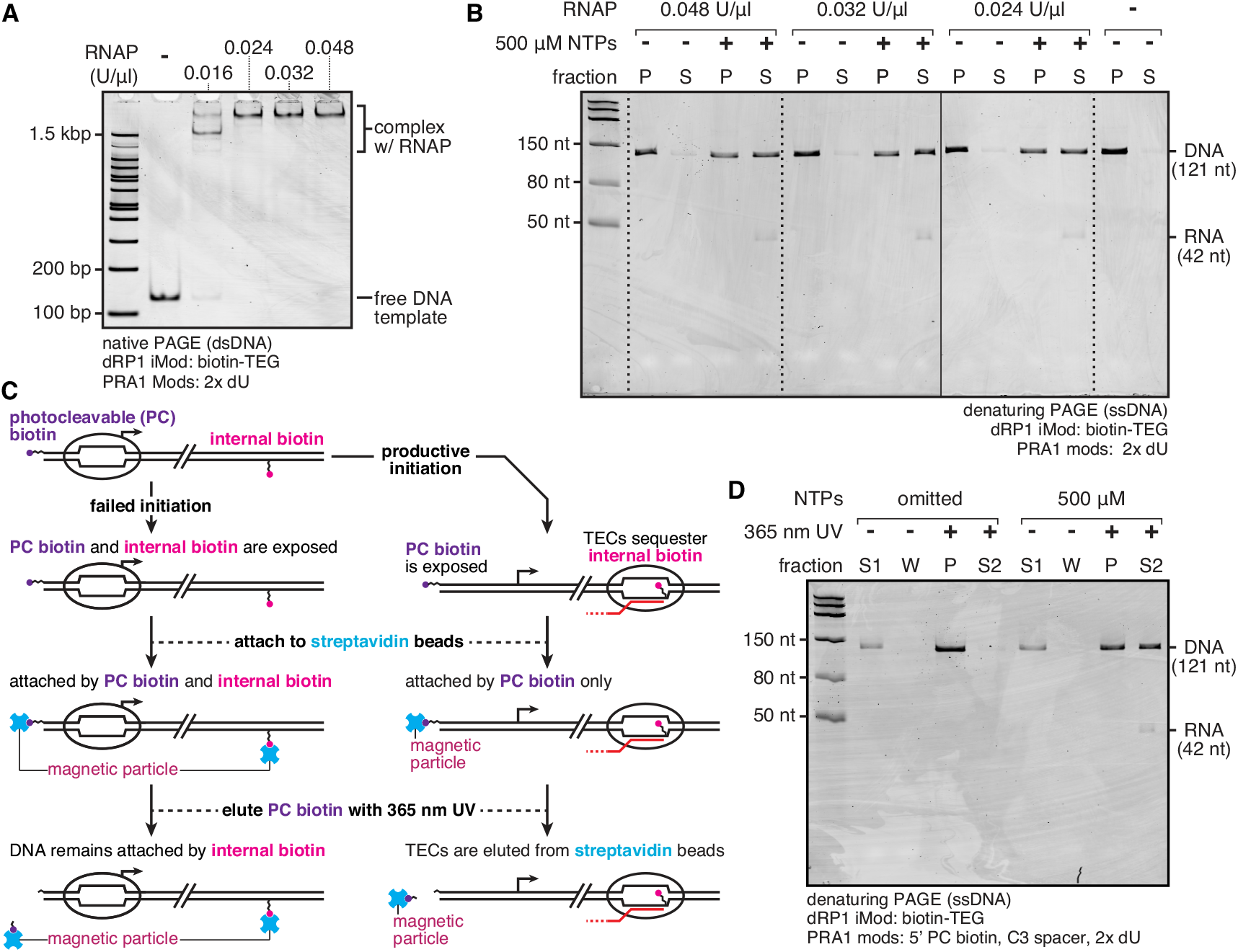
Selective elution of roadblocked TECs from streptavidin beads. **(A)** EMSA of open complexes formed with 5 nM DNA template and variable concentrations of *E. coli* RNAP. **(B)** Fractionation of internal biotin-TEG-modified DNA template following transcription with variable concentrations of *E. coli* RNAP in the presence and absence of NTPs. P = Pellet, S = Supernatant. The solid vertical line between 0.032 and 0.024 U/μl samples indicates a gel splice. **(C)** Overview of a strategy for purification of TECs. If RNAP fails to escape the promoter, DNA is attached to streptavidin beads by both PC biotin and internal biotin-TEG. If RNAP escapes the promoter, internal biotin-TEG is sequestered by a TEC, so that DNA is only attached to streptavidin beads by PC biotin. TECs can therefore be selectively photo-eluted by 365 nm UV. **(D)** Proof-of-principle of the TEC purification procedure shown in (C). S1 is the supernatant after bead binding, W is the supernatant after a wash step to remove transcription reaction components, P is the pellet, and S2 is the supernatant following photo-elution. The experiment in (A) was performed once to set conditions for (B) and (D). The gels in (B) and (D) are representative of two independent replicates.

TEC-dependent exclusion from streptavidin beads provides a basis for the purification of TECs but must be coupled with a gentle buffer exchange procedure so that components of the transcription reaction can be depleted prior to downstream assays. To this end, I designed the following purification strategy (Fig. 3C): In addition to the internal biotin-stall site, a 5’ PC biotin modification (38) is included in the DNA template (DNA Template 4 in Table S2). If RNAP fails to escape the promoter, the DNA is attached to a magnetic particle both by the internal biotin and the PC biotin; bead immobilization is performed under dilute conditions to maximize the likelihood that both biotin modifications are attached to the same bead (Experimental Procedures). In contrast, if RNAP escapes the promoter the internal biotin modification is sequestered by RNAP so that the DNA is attached to the beads by the PC biotin alone. Consequently, upon irradiation by 365 nm UV LEDs (Fig. S1), DNA containing a TEC at the internal biotin stall site is eluted into the supernatant and DNA that does not contain a stalled TEC is retained in the bead pellet.

To establish a proof-of-principle for this TEC purification strategy, I used 5 nM DNA template and 0.024 U/μl RNAP in a 25 μl reaction in order to saturate transcription (Fig. 3A); further optimizations are described in the next Results section. The purification was performed as follows (see Experimental Procedures for complete details): After forming open promoter complexes in the absence or presence of 500 μM NTPs, single-round transcription was initiated by the simultaneous addition of MgCl_2_ to 10 mM and rifampicin. After two minutes of transcription, the reaction was diluted by adding 200 μl of Transcription Buffer supplemented with rifampicin and mixed with 25 μl of 1 μg/μl streptavidin beads in Transcription Buffer. This bead-binding mixture was incubated for 1 hour at room temperature with rotation, the beads were pelleted using a magnet stand, and the supernatant was collected as fraction S1. The beads were resuspended in 250 μl of Transcription Buffer supplemented with 1 mM MgCl_2_, pelleted again using a magnet stand, and the wash supernatant was collected as fraction W. The beads were then resuspended in 25 μl of Transcription Buffer supplemented with 1 mM MgCl_2_ and irradiated with 365 nM UV LEDs (∼10 mW/cm^2^ from four directions) for 5 minutes. The bead pellet was collected as fraction P and the supernatant as fraction S2. In both the absence and presence of NTPs ∼17% of DNA was lost in fraction S1, presumably because the reaction was diluted for bead binding to avoid crosslinking the streptavidin beads with the doubly biotinylated DNA template (Fig. 3D). No nucleic acids were detectable in fraction W, indicating that the buffer exchange step does not incur substantial sample loss (Fig. 3D). In the absence of NTPs, ∼2% of the remaining DNA was present in the bead pellet following UV irradiation, indicating that the majority of DNA is retained on the beads by the internal biotin modification (Fig. 3D). In contrast, when TECs were formed by the inclusion of NTPs, ∼46% of the remaining DNA was present in fraction S2 (Fig. 3D). Consistent with the interpretation that the DNA collected in fraction S2 in the presence of NTPs is due to TEC-dependent streptavidin bead exclusion, the 42 nt RNA transcript was only detectable in fraction S2 (Fig. 3D).

### A DNA competitor strategy excludes open complexes from purified TEC DNA

The proof-of-principle TEC purification shown in Figure 3D used a saturating RNAP concentration to assess the efficiency limit of the preparation. However, because this procedure uses rifampicin to limit transcription to a single round (as opposed to heparin, which could interfere with downstream assays (39)), these conditions are likely non-optimal for the preparation of pure TECs: after promoter escape, free RNAP holoenzyme can bind unoccupied promoters to yield DNA that contains both a TEC and an open complex. Consistent with this expectation, EMSAs of prepared TECs revealed the presence of two bands, suggesting that the preparations contain two distinct species (Fig. 4B, lanes 2 and 3). In support of an interpretation that the faster migrating band corresponds to pure TECs and the slower migrating band corresponds to TECs on DNA with an associated open complex, decreasing the ratio of RNAP to DNA template so that open complex formation was not saturated caused the faster migrating band to become more prominent(Figs. 4A, 4B (lanes 2, 3, and 4)).

**Figure 4.**
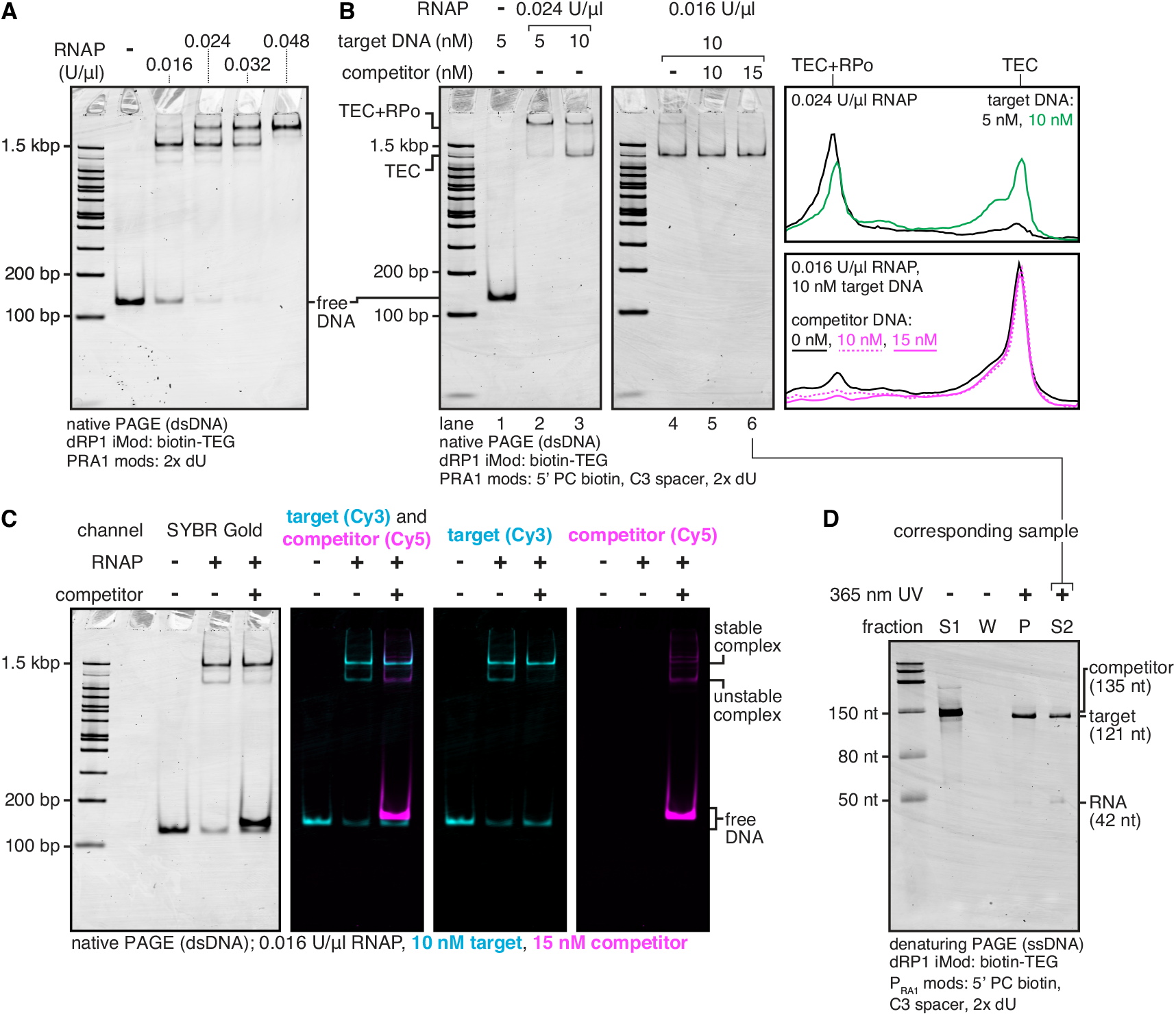
Optimization of TEC purification. **(A)** EMSA of open complexes formed with 10 nM DNA template and variable concentrations of *E. coli* RNAP. **(B)** EMSA of TECs purified with variable DNA template, RNAP, and competitor DNA template concentrations. The assay shown in Figure 5 revealed the slow-migrating band to be TECs with an associated open promoter complex, and the fast-migrating band to be pure TECs. Independent TEC preparations using the 0.016 U/μl RNAP, 10 nM target DNA, and 15 nM competitor DNA condition from lane 6 are shown in Figures S2, S3A, and S3B. **(C)** EMSA of sequentially formed of open complexes with Cy3-labelled target and Cy5-labelled competitor DNA. **(D)** Denaturing PAGE of purification fractions for TECs prepared using the conditions in (B) lane 6. S1 is the supernatant after bead binding, W is the supernatant after a wash step to remove transcription reaction components, P is the pellet, and S2 is the supernatant following photo-elution. The experiment in (A) was performed once to set conditions for (B), (C), and (D). The gels in (B), (C), and (D) are representative of two independent replicates.

Reactions that used 10 nM DNA template and 0.016 U/μl RNAP nearly eliminated the slower migrating band (Fig. 4B, lane 4) (note that when preparing TECs with 10 nM DNA, the amount of streptavidin beads was doubled, and the binding reaction diluted to 500 μl accordingly. See Experimental Procedures).

Following the observation that decreasing the RNAP:DNA ratio increased the homogeneity of TEC preparations, I implemented a DNA competitor strategy to scavenge free RNAP holoenzyme before initiating transcription. Below, ‘Target DNA’ refers to the doubly biotinylated DNA template that is the intended substrate for TEC purification (DNA Template 4 in Table S2) and ‘Competitor DNA’ refers to the DNA template used to scavenge free RNAP (DNA Template 5 in Table S2). In this strategy, open complexes are first formed on the Target DNA using 10 nM DNA template and 0.016 U/μl RNAP, which alone yielded a fast-migrating product with ∼94% purity (Fig. 4B, lane 4). The transcription reaction is then mixed with Competitor DNA containing the P_RA1_ promoter to scavenge free RNAP in open complexes, and an internal etheno-dA stall site to retain RNAP on the Competitor template if promoter escape succeeds. Visualization of this procedure using Cy3-labelled Target DNA and Cy5-labelled Competitor DNA (DNA Templates 6 and 7 in Table S2, respectively) revealed two populations of Target DNA:RNAP complexes: stable complexes that resisted the Competitor DNA and a distinct population of unstable complexes that were titrated to Competitor DNA (Fig. 4C). The presence of substantial free Competitor DNA after open complex formation suggests that most RNAP is engaged in a complex with DNA (Fig. 4C). As expected, scavenging free RNAP prior to TEC purification further reduced the slow-migrating product so that, in reactions that contained 10 nM Target DNA and 0.016 U/μl RNAP, the fast-migrating product was consistently >97% pure (Figs. 4B (lanes 5 and 6), and S2). Competitor DNA was efficiently removed from the TEC preparations in the S1 fraction (Fig. 4D). It was not possible to determine the fraction of input Target DNA that was present in S1 because Target and Competitor DNA differ in length by only 14 nt, however it is clearly a small fraction relative to bead-bound DNA (Fig. 4D). Of the DNA that bound to the streptavidin beads, ∼37% was released into fraction S2 after photo-elution (Fig. 4D).

To verify that the fast- and slow-migrating products observed by EMSA correspond to pure TECs and TECs with an associated open complex, respectively, I performed a ‘USER^®^ footprinting assay’ to directly assess promoter accessibility in TECs prepared under different conditions. Because the dU bases in the P_RA1_ promoter are in close contact with σ^70^, open promoter complexes interfere with USER^®^ digestion even when long (30 min at 37°C) incubation times are used (Fig. 5B, C; compare lanes 2 and 3). When TECs were prepared under conditions that primarily yielded the slow-migrating product, USER^®^ digestion was severely attenuated (Fig. 5B, C; lane 5). In contrast, digestion of TECs that were prepared under conditions that yield a >97% pure fast-migrating product were digested as effectively as naked DNA (Fig. 5B, C; lane 7).

**Figure 5.**
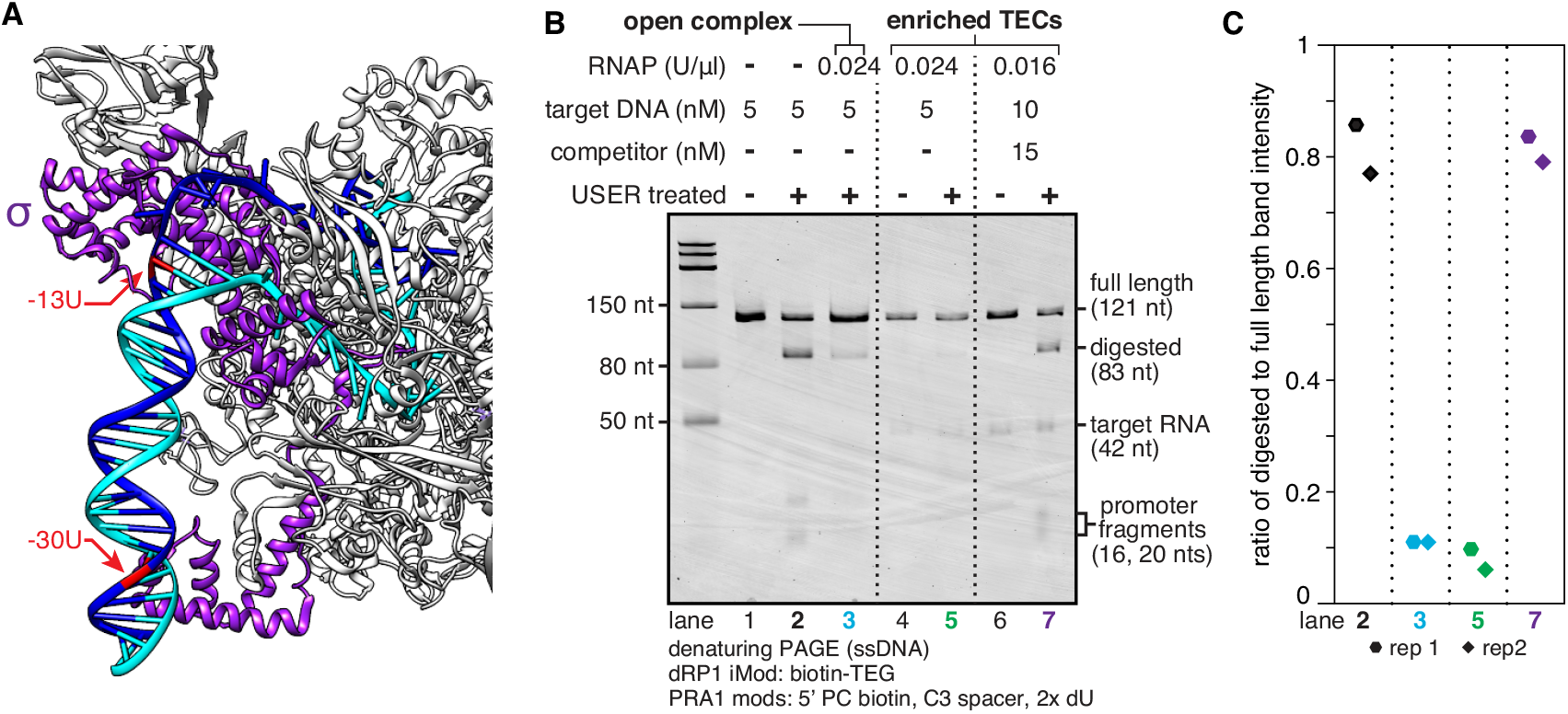
USER^®^ footprinting to assess promoter accessibility. **(A)** Structure of *Thermus aquaticus* open promoter complex (PDB:4XLN) (50) annotated to indicate the position of P_RA1_ dU bases relative to direct contacts between σ and promoter DNA. **(B)** USER^®^ footprinting assay to assess P_RA1_ promoter accessibility. Saturation of the P_RA1_ promoter with open complexes (conditions from Figure 3A) attenuates USER^®^ digestion. TECs prepared under conditions that predominantly yield slow-migrating complexes (Figure 4B, lane 2) inhibit USER^®^ digestion, indicating promoter occupation by open complexes. TECs prepared under conditions that yield >97% fast-migrating complexes (Figure 4B, lane 6) are efficiently digested, indicating the absence of open complexes. **(C)** Quantification USER^®^ digestion for key lanes from (B) as the ratio of USER^®^ digested to full length band intensity. The gel in (B) is representative of two independent replicates.

### Purified TECs are compatible with P_RA1_ barcoding

TEC purification and P_RA1_ barcoding both depend on the internal biotin-TEG modification: during TEC purification, the internal biotin-TEG modification is used both as an RNAP stall site and as an attachment point for depleting DNA that does not contain a TEC (Fig. 3C). During P_RA1_ barcoding, the internal biotin modification must be available as an attachment point for depleting excess barcoding oligonucleotide (Fig. 2B). Consequently, compatibility of the TEC purification and P_RA1_ barcoding protocols requires efficient TEC degradation. Most TECs were completely dissociated from DNA following treatment with Thermolabile Proteinase K and trace amounts of remaining TECs were dissociated after heating to 65°C for 10 minutes (Fig. S3A). Similarly, the RNA transcript was readily degraded by treatment with RNase I_f_, which was heat-inactivated by heating to 70°C for 20 minutes (Fig. S3B). The resulting DNA was then readily processed by the P_RA1_ barcoding protocol and amplified using a standard Illumina RNA PCR Index Primer and an extended version of the Illumina RNA PCR Primer (Table S1), indicating that TEC purification and P_RA1_ barcoding can be integrated using only enzymatic reactions and heat inactivation steps (Fig. S3B).

## Discussion

I have developed an integrated procedure that combines purification of precisely positioned *E. coli* RNAP TECs with quantitative DNA barcoding. This TEC Display method establishes a platform for the development of multiplexed assays in which a TEC library is partitioned by its functional properties so that the activity of sequence variants can be inferred from their distribution across fractions.

In the present study, I have established procedures for the front (TEC purification) and back (quantitative DNA barcoding) ends of TEC Display experiments. Application of TEC Display to address biological questions will depend on the development of ‘fractionation-by-function’ approaches for assessing RNA activity. In principle, the TEC Display platform described here could be applied to study the basis of interactions between RNA and other biomolecules such as proteins (40), small molecules (41), and other RNAs (42). Neither the concept of assessing RNA function using cotranscriptionally displayed sequence libraries nor the concept of fractionating an RNA library by its affinity for an immobilized target are new: RNA-HiTS experiments assess RNA function by cotranscriptionally displaying RNA libraries *in situ* on an Illumina sequencer flow cell for fluorescence-based assays (6,7), and RNA Bind-n-Seq (RBNS) quantifies RNA-binding protein specificity by fractionating an RNA library based on its affinity for a target protein (43). The central advance of the work presented here is that, by integrating TEC purification with quantitative DNA barcoding, TEC Display combines features of its predecessors in a way that will enable assays that use cotranscriptionally displayed RNA libraries to be performed at the laboratory bench.

TEC Display was designed with an emphasis on broad accessibility, the potential for scalability, and versatility. TEC purification and DNA barcoding can be performed with commercially available reagents and, with the exception of a custom-built 365 nm UV microcentrifuge tube irradiator (Fig. S1), common laboratory equipment. It should therefore be possible for any laboratory with biochemistry or molecular biology expertise to use or develop TEC Display assays. Towards the goal of scalability, TEC Display makes extensive use of magnetic bead immobilization and thermolabile enzymes to circumvent the need for organic extractions or precipitations. It is therefore likely that TEC Display procedures can be automated. Furthermore, TEC display is immediately compatible with cotranscriptional RNA structure probing methods (9,44) and can likely be integrated into *in vitro* selection workflows (22,23). For these reasons I anticipate that, through its application and continued technical expansion, TEC Display will complement existing approaches for systematic analysis of cotranscriptionally displayed RNA.

The experimental procedures described here are enabled by a method I previously developed for the sequence-independent preparation of internally modified dsDNA, which can be used to halt RNAP one nucleotide upstream of a functionalized chemical lesion (24). TEC Display provides the first example of how this chemical transcription roadblocking approach can facilitate intricate manipulations of macromolecular complexes: In the combined TEC purification and DNA barcoding procedure, the internal biotin-TEG modification functions first as a transcription stall site, then as a means for selecting TECs, and finally as an attachment point for purifying the barcoded DNA library away from excess reaction components. Furthermore, an etheno-dA stall site (24,45) in the competitor DNA used to scavenge free RNAP during open complex formation is used to trap any RNAP that escapes the promoter upon transcription initiation. Currently four DNA modifications have been validated for transcription roadblocking using internally modified DNA, and extension of this approach to include other DNA modifications with distinct functions will further expand our ability to manipulate TECs in complex procedures.

Beyond its immediate applications for TEC Display, my procedure for purifying TECs by selective photo-elution may be generally useful for studies that involve assembling transcription complexes *in vitro*. Structural and biochemical investigations that require the assembly of defined TECs have classically used a ‘scaffold’ approach in which ternary complexes are assembled by the sequential addition of nucleic acid and protein components (46,47). For many applications this approach is optimal, however some biologically important transcription complexes may require co-transcriptional or promoter-directed assembly. In such cases, my selective photo-elution method provides a straightforward strategy for bead-based preparation of defined TECs following promoter-directed transcription initiation. Furthermore, while the procedures I have presented here focus on the purification of TECs, it is likely that different DNA modification configurations can facilitate the preparation of other transcription complexes.

### Experimental Procedures

#### Oligonucleotides

All oligonucleotides were purchased from Integrated DNA Technologies (IDT). A detailed description of all oligonucleotides including sequence, modifications, and purifications is presented in Table S1. Oligonucleotides and DNA preparations that contained a 5’ PC biotin modification were handled under low intensity 592 nm light from SimpleColor™ Amber LEDs (Waveform Lighting) out of an abundance of caution and stored as single-use aliquots. Cy3- and Cy5-labelled oligonucleotides and DNA preparations were handled under low intensity room light and stored as single-use aliquots.

#### Proteins

All enzymes, including Q5^®^ High-Fidelity DNA Polymerase, Q5U^®^ High-Fidelity DNA Polymerase, Vent^®^ (exo-) DNA polymerase, *Sulfolobus* DNA Polymerase IV, Thermolabile Exonuclease I, Klenow Fragment (3’→5’ exo-), Klenow Fragment, T4 DNA Polymerase, Thermolabile USER^®^ II Enzyme, T4 DNA Ligase, *E. coli* RNA Polymerase holoenzyme, Thermolabile Proteinase K, and RNase I_f_ were purchased from New England Biolabs (NEB).

#### Sequences

Annotated sequences of the constant regions required for TEC Display can be viewed at https://benchling.com/s/seq-DOgBWf14L74Gd9rVv7nc. The sequence of a TEC Display library after barcoding and PCR amplification with full annotations can be viewed at https://benchling.com/s/seq-QAf9ODTHtJT1S7aNJ49R. Illumina adapter sequences are from the TrueSeq Small RNA kit. In the sequence annotations, VRA3 is the reverse complement of the Illumina RA5 adapter, and VRA5 is the reverse complement of the Illumina RA3 adapter; this notation was originally used in the Precision Run-On Sequencing method (48,49).

#### DNA template preparation

Several DNA template preparation protocols were performed depending on the requirements of oligonucleotide modifications. For simplicity, each possible processing step is detailed below in general terms, and Table S2 provides details on the oligonucleotides and specific processing steps used for every DNA template preparation in this work. All DNA templates that required translesion synthesis were assessed by both denaturing and non-denaturing PAGE quality control analyses, which are shown in Figure S4.

##### PCR Amplification

DNA templates with dU bases were prepared using Q5U^®^ High-Fidelity DNA Polymerase (NEB) in reactions that contained 1X Q5U^®^ Buffer (NEB), 0.2 mM dNTPs (Invitrogen), 0.25 μM forward primer, 0.25 μM reverse primer, 20 pM template oligonucleotide, and 0.02 U/μl Q5U^®^. Details of the oligonucleotides used for each DNA template preparation are available in Tables S1 and S2. DNA templates that did not contain dU bases were prepared in a reaction that used Q5^®^ High-Fidelity DNA Polymerase (NEB) but was otherwise identical. Typically, six 100 μl reaction volumes were prepared on ice and aliquoted into 200 μl thin-walled tubes. The thermal cycling protocol was: 98 °C for 30 s, [98 °C for 10 s, 65 °C for 20 s, 72 °C for 20 s] x 30 cycles, 72 °C for 5 minutes, hold at 12 °C.

##### Translesion Synthesis

After PCR amplification, DNA templates that required translesion synthesis were purified using a QIAquick^®^ PCR Purification Kit (Qiagen) (1 column per 100 μl reaction) and eluted in 50 μl of Buffer EB (10 mM Tris pH 8.5); one negative control reaction was eluted in 30 μl of Buffer EB and not processed further. This PCR purification step is required to exchange the reaction buffer and deplete dNTPs. 500 μl translesion synthesis reactions contained 1X ThermoPol^®^ Buffer (NEB), 200 μM dGTP (Invitrogen), 200 μM dTTP (Invitrogen), 200 μM 2-Amino-dATP (TriLink Biotechnologies), 200 μM 5-Propynyl-dCTP (TriLink Biotechnologies), 0.02 U/μl Vent^®^ (exo-) DNA Polymerase, and 0.02 U/μl *Sulfolobus* DNA Polymerase IV and were split into 100 μl reaction volumes in 200 μl thin-walled tubes. Translesion synthesis reactions were incubated at 55 °C in a thermal cycler with a heated lid set to 105 °C for 1 hour.

##### DNA purification by agarose gel extraction

Agarose gel purification was performed essentially as described previously (24). Five 100 μl PCR or translesion synthesis reactions were pooled and ethanol precipitated by adding 50 μl of 3M sodium acetate (NaOAc) (pH 5.5) and 1 ml of cold 100% ethanol and chilled at -70 °C for 30 minutes before centrifugation at 18,500 x g and 4 °C for 30 minutes. Pelleted DNA was washed with 1 ml of cold 70% ethanol, air-dried, and resuspended in 30 μl of 10 mM Tris-HCl (pH 8.0). Samples were mixed with 6X XC Only SDS DNA Loading Dye (30% glycerol, 10 mM Tris-HCl (pH 8.0), 0.48% (w/v) sodium dodecyl sulfate (SDS), 0.01% xylene cyanole FF), run on a tris-borate-EDTA (TBE) 1% (wt/v) agarose gel that was prepared using SeaKem^®^ GTG Agarose (Lonza Bioscience) and contained 0.625 μg/ml ethidium bromide. All DNA was able to be excised without UV exposure because the co-migration of ethidium bromide made the large quantity of DNA readily observable under visible light. DNA was purified using a QIAquick^®^ Gel Extraction Kit (Qiagen) according to the manufacturer’s protocol, except that agarose gel slices were melted at 30 °C and the DNA was eluted in 30 μl of 10 mM Tris-HCl (pH 8.0). DNA concentration was quantified using a Qubit™ 4.0 Fluorometer (Invitrogen) with the Qubit™ dsDNA Broad Range Assay Kit (Invitrogen) according to the manufacturer’s protocol.

##### DNA clean-up by exonuclease digestion

All DNA containing light-sensitive modifications was purified using an alternate protocol that used exonuclease digestion to degrade trace amounts of excess primers in place of gel extraction to minimize light exposure. This procedure was possible because all DNA was prepared using an oligonucleotide as the PCR template, so it was not necessary to remove plasmid DNA. PCR or translesion synthesis reactions were pooled and 0.5 μl of Thermolabile Exonuclease I (NEB) was added per 100 μl reaction volume. Reactions were incubated 37 °C for 4 minutes using a thermal cycler and placed on ice. The thermal cycler block was set to 80°C and, after it reached temperature, the reactions were returned to the block for 1 minute to heat inactivate the Thermolabile Exonuclease I. The reactions were then purified using a QIAquick PCR Purification Kit (up to 2.5 reactions per column) following the manufacturer’s protocol, except that two 750 μl Buffer PE washes were performed, and the samples were eluted in 50 μl of 10 mM Tris-HCl (pH 8.0). The concentration of purified DNA was quantified using a Qubit™ 4.0 Fluorometer with the Qubit™ dsDNA Broad Range Assay Kit according to the manufacturer’s protocol. It is important to note that the Thermolabile Exonuclease I digestion was able to be performed immediately following translesion synthesis because primers were depleted to undetectable levels during the Q5U^®^ and Q5^®^ PCRs (Fig. S4). My laboratory has observed that under PCR amplification conditions that do not lead to virtually complete primer depletion, the presence of excess oligonucleotides and dNTPs can lead to the formation of primer dimers because *Sulfolobus* DNA polymerase IV is active at 37 °C, which permits the formation of primer dimer products that do not form at higher temperatures. In DNA preparations where primers are not depleted during PCR, this issue is resolved simply by performing an additional PCR clean-up immediately after the translesion synthesis reaction to remove the DNA polymerases and free dNTPs.

### Preparation of streptavidin-coated magnetic beads for P_RA1_ barcoding assays

For P_RA1_ barcoding assays 10 μl of Dynabeads™ MyOne™ Streptavidin C1 beads (Invitrogen) per 25 μl sample volume were prepared in bulk: After placing the beads on a magnet stand and removing the storage buffer, the beads were resuspended in 500 μl of Hydrolysis Buffer (100 mM NaOH, 50 mM NaCl) and incubated at room temperature for 10 minutes with rotation. Hydrolysis Buffer was removed, and the beads were resuspended in 1 ml High Salt Wash Buffer (50 mM Tris-HCl (pH 7.5), 2 M NaCl, 0.5% Triton X-100), transferred to a new tube, and washed by rotating for 5 minutes at room temperature. High Salt Wash Buffer was removed, and the beads were resuspended in 1 ml Binding Buffer (10 mM Tris-HCl (pH 7.5), 300 mM NaCl, 0.1 % Triton X-100), transferred to a new tube, and washed by rotating for 5 minutes at room temperature. Binding buffer was removed, and the beads were resuspended in 25 μl of per sample of Binding Buffer and transferred to a new tube. Beads were prepared fresh for each experiment and kept on ice until use.

### P_RA1_ barcoding validation assays

The barcoding reaction validation assays in Figure 2 were performed using a single master mix from which fractions were taken during processing to visualize intermediate steps. The volume of an individual sample within the master mix was 25 μl. For clarity and simplicity, the procedure below specifies the volume of each reagent added to the master mix per 25 μl sample volume. A description of the intermediate fractions that were collected is provided at the end of this sub-section.

The barcoding reaction master mix contained 1X T4 DNA Ligase Buffer (NEB) (2.5 μl/sample volume of 10X T4 DNA Ligase Buffer), 5 nM DNA template (0.125 pmol/sample volume), and 0.02 U/μl Thermolabile USER^®^ II Enzyme (NEB) (0.5 μl/sample volume) and was prepared on ice in a thin-walled 200 μl tube. The dU excision reaction was incubated at 37 °C for 30 min on a thermal cycler block with a heated lid set to 45 °C. After dU excision, 0.5 μl/sample volume of 2.5 μM barcoding oligo PRA1m12_VRA5_16N (Table S1) was added to the master mix. To anneal the barcoding oligo and inactivate Thermolabile USER^®^ II Enzyme, the master mix was placed on a thermal cycler block set to 70 °C with a heated lid set to 105 °C and slowly cooled using the protocol: 70 °C for 5 minutes, ramp to 65 °C at 0.1 °C/s, 65 °C for 5 minutes, ramp to 60 °C at 0.1 °C/s, 60 °C for 2 minutes, ramp to 25 °C at 0.1 °C/s, hold at 25 °C. After annealing the barcoding oligo, 1 μl/sample volume of T4 DNA ligase (NEB) was added to the master mix. The ligation reaction was incubated at 25 °C for 1 hour, and T4 DNA ligase was inactivated by incubation at 65 °C for 10 minutes.

After the ligation step was completed, the master mix was transferred to a 1.7 ml microcentrifuge tube and mixed with 26.5 μl/sample volume of 2X Binding Buffer (20 mM Tris-HCl, (pH 7.5), 600 mM NaCl, 0.2% Triton X-100). Pre-equilibrated streptavidin-coated magnetic beads were placed on a magnet stand and the Binding Buffer used for bead storage was removed. The beads were resuspended with the master mix and incubated at room temperature with rotation for 30 minutes. After bead binding, the sample was briefly spun down in a mini centrifuge and placed on a magnet stand for 1 minute. The supernatant was removed, and the sample was briefly spun down a second time and returned to the magnet stand for removal of residual supernatant. The sample was then resuspended in 100 μl/sample volume of 1X NEBuffer 2 (NEB) supplemented with Triton X-100 to 0.1% and transferred to a new tube. If the master mix still contained multiple sample volumes at this point, it was divided into individual 100 μl samples. Samples were incubated at room temperature with rotation for 5 minutes, briefly spun down, and placed on a magnet stand for 1 minute. The supernatant was removed, and the samples were briefly spun down a second time and returned to the magnet stand to remove residual supernatant. The beads were then resuspended in 100 μl of Extension Master Mix, which varied by experiment. 100 μl extension reactions were performed with Klenow Fragment (3’→5’ exo-) (NEB) (Figs. 2C, 2E), Klenow Fragment (NEB) (Fig. 2E, 2F), or T4 DNA Polymerase (NEB) (Fig. 2E). Klenow Fragment (3’→5’ exo-) reactions contained 1X NEBuffer 2, 0.2 mM dNTPs, and 0.1 U/μl Klenow Fragment (3’→5’ exo-) and were incubated at 25 °C for 15 minutes followed by 37 °C for 15 minutes. Klenow Fragment reactions contained 1X NEBuffer 2, 0.2 mM dNTPs, and 0.05 U/μl Klenow Fragment, and were incubated at 25 °C for 20 minutes. T4 DNA Polymerase reactions contained 1X NEBuffer 2.1 (NEB), 0.2 mM dNTPs and 0.03 U/μl T4 DNA Polymerase and were incubated at 12 °C for 20 minutes. Following extension, the reactions were transferred to a pre-chilled 1.7 ml microcentrifuge tube, placed on a magnet stand, and the supernatant was removed. The barcoded DNA was then eluted from the beads using either a denaturing or non-denaturing procedure: DNA containing either a 5’ or internal biotin modification was recovered by resuspending the bead pellet in 25 μl of 95% formamide and 10 mM EDTA, heating at 100 °C for five minutes, placing the sample on a magnet stand, and collecting the supernatant. DNA containing an internal desthiobiotin was recovered by resuspending the bead pellet in 150 μl of Biotin Elution Buffer (0.5 M Tris-HCl (pH 8.0), 10 mM EDTA (pH 8.0), 10% DMSO, 2 mM D-Biotin), incubating at room temperature with rotation for 15 minutes, placing the sample on a magnet stand, and collecting the supernatant.

Intermediate fractions were collected during the barcoding protocol as follows: Unprocessed control fractions were collected by transferring 24.5 μl of the initial master mix to 125 μl of Stop Solution (0.6 M Tris-HCl (pH 8.0), 12 mM EDTA (pH 8.0)) before Thermolabile USER^®^ II Enzyme was added. USER^®^ digested fractions were collected by transferring 25 μl of master mix to 125 μl of Stop Solution after the dU excision reaction. Barcode ligated fractions were collected by transferring 26.5 μl of master mix to 125 μl of Stop Solution after T4 DNA ligase was inactivated. Wash fractions were collected by transferring a wash supernatant volume that corresponded to one sample to Stop Solution for a total volume of 150 μl. Fully barcoded fractions that were eluted from beads by heat denaturation were collected by transferring the 25 μl supernatant to 125 μl of Stop Solution. Fully barcoded fractions that were eluted in 150 μl of Biotin Elution Buffer were collected directly.

All fractions were processed as follows: 150 μl of Tris (pH 8) buffered phenol:chloroform:isoamyl alcohol (25:24:1, v/v) (Thermo Scientific) was added to each sample. Samples were mixed by vortexing and inversion and centrifuged at 18,500 x g and 4 °C for five minutes. The aqueous phase was collected and transferred to a new tube. DNA was precipitated by adding 15 μl 3 M NaOAc (pH 5.5), 450 μl 100% ethanol, and 1.5 μl GlycoBlue™ Coprecipitant (Invitrogen) to each sample. The samples were chilled at -70 °C for 30 minutes and centrifuged at 18,500 x g and 4 °C for 30 minutes. Samples for denaturing PAGE were resuspended in 32 μl of Formamide Loading Dye (90% (v/v) deionized formamide, 1X Transcription Buffer (see below), 0.025% (w/v) bromophenol blue, 0.025% (w/v) xylene cyanole FF); half the sample was used for denaturing PAGE. Samples for native PAGE were resuspended in 10 μl of 10 mM Tris-HCl (pH 8.0) and 2 μl of 6X DNA loading dye (30% (v/v) glycerol, 0.025% (w/v) bromophenol blue, 0.025% (w/v) xylene cyanole FF).

### Denaturing Urea-PAGE

Gels for Urea-PAGE were prepared at 10% or 12% using the SequaGel UreaGel 19:1 Denaturing Gel System (National Diagnostics). The conditions used for denaturing PAGE were selected to ensure complete and consistent denaturation of dsDNA using a Mini-PROTEAN^®^ Tetra Vertical Electrophoresis Cell (Bio-Rad): The inner buffer chamber was filled completely with 1X TBE, but the outer buffer chamber contained enough 1X TBE to cover only ∼1 cm of the gel plates to reduce heat loss. Urea-PAGE gels were run at 480 V, which is 80% of the maximum voltage recommended for the Mini-PROTEAN^®^ system. Urea-PAGE gels were stained with 1X SYBR Gold (Invitrogen) in 1X TBE for 10 minutes and scanned on an Sapphire™ Biomolecular Imager (Azure Biosystems) using the 488 nm/518BP22 setting.

### Native PAGE for the analysis of purified DNA

Purified DNA was assessed using 1X TBE 8% polyacrylamide gels prepared using ProtoGel (National Diagnostics) acrylamide. Native PAGE gels were stained with 1X SYBR Gold in 1X TBE for 10 minutes and scanned on a Sapphire™ Biomolecular Imager using the 488 nm/518BP22 setting.

### Analysis of open complex formation by EMSA

For Figures S3A and 4A, reactions containing 1X Transcription Buffer (20 mM Tris-HCl (pH 8.0), 50 mM KCl, 1 mM DTT, 0.1 mM EDTA (pH 8.0)), 0.1 mg/ml BSA (Invitrogen), 5 nM (Fig. 3A) or 10 nM (Fig. 4A) DNA Template 2 (Table S2), and 0, 0.016, 0.024, 0.032, or 0.048 U/μl *E. coli* RNAP holoenzyme (NEB) were incubated in a dry bath set to 37 °C for 15 (Fig. 3A) or 20 (Fig. 4A) minutes. After incubation, 15 μl of sample was mixed with 3 μl of 6X BPB Only Native DNA Loading Dye (30% (v/v) glycerol, 10 mM Tris-HCl (pH 8.0), 0.01% (w/v) bromophenol blue), loaded on a 0.5X TBE 5% polyacrylamide gel that was submerged up to the wells in 0.5X TBE, and run at 45 V for ∼1.5 hr. Gels were stained with 1X SYBR Gold in 0.5X TBE for 10 minutes and scanned on a Sapphire™ Biomolecular Imager using the 488 nm/518BP22 setting.

For Figure 4C, open complexes were prepared as described below in the section *Purification of TECs by selective photo-elution from streptavidin beads* except that NTPs were omitted from the reaction, DNA Templates 6 and 7 (Table S2) were used as the ‘Target’ and ‘Competitor’ respectively, and the reactions were performed under low intensity room light instead of 592 nm light. Reactions were staggered so that all sample handling reached completion simultaneously. 15 μl of sample was mixed with 3 μl of 6X BPB Only Native DNA Loading Dye and loaded on a 0.5X TBE 5% polyacrylamide gel as described above except that the gel was run in a dark room. After the gel had run, it was transferred to a plastic dish containing 0.5X TBE, scanned sequentially on a Sapphire™ Biomolecular Imager using the 520 nm/565BP24 and 658 nm/710BP40 settings, removed from the imager, stained with 1X SYBR Gold in 0.5X TBE for 10 minutes and scanned again on a Sapphire™ Biomolecular Imager using the 488 nm/518BP22 setting.

### Internal biotin-TEG sequestration assay

10 μl of Dynabeads™ MyOne™ Streptavidin C1 beads per 25 μl sample volume were prepared in bulk as described above in *Preparation of streptavidin-coated magnetic beads for P*_*RA1*_ *barcoding assays*, except that two additional washes were performed using 500 μl of Wash Buffer T (1X Transcription Buffer supplemented with 0.1% Triton X-100) and each sample volume of beads was stored in a 25 μl aliquot in Wash Buffer T.

All 25 μl transcription reactions contained 1X Transcription Buffer, 0.1 mg/ml BSA, and 5 nM DNA Template 2 (Table S2). NTPs (GE Healthcare) were either omitted or included at 500 μM each; *E. coli* RNAP holoenzyme was either omitted or included at 0.024, 0.032, or 0.048 U/μl. At the time of preparation the total reaction volume was 22.5 μl due to the omission of 10X Start Solution (100 mM MgCl_2_, 100 μg/ml rifampicin).

Reactions were placed in a dry bath set to 37 °C for 15 minutes to form open promoter complexes. 2.5 μl of 10X Start Solution was added and transcription was allowed to proceed for 2 minutes during which an aliquot of beads was placed on a magnet stand to remove the Wash Buffer T. After 2 minutes of transcription, the pelleted beads were resuspended using the 25 μl transcription reaction, placed on a rotator, and incubated at room temperature for 30 minutes. The beads were then returned to the magnet stand and the supernatant was separated from the pellet and added to 125 μl of Stop Solution. To recover immobilized DNA, the bead pellet was resuspended in 25 μl of 95% formamide and 10 mM EDTA, heated at 100 °C for five minutes, placed on a magnet stand, and the supernatant was collected and added to 125 μl of Stop Solution. Reactions were processed for denaturing Urea-PAGE by phenol:chloroform extraction and ethanol precipitation as described above in the section *P*_*RA1*_ *barcoding validation assays* and resuspended in 16 μl of BPB Only Formamide Loading Dye (90% (v/v) deionized formamide, 1X Transcription Buffer, 0.01% (w/v) bromophenol blue). Urea-Page was performed as described above in the section *Denaturing Urea-PAGE*.

### Preparation of streptavidin beads for TEC Purification

For TEC purification assays, 2.5 μl or 5 μl of Dynabeads™ MyOne™ Streptavidin C1 beads per 25 μl sample volume were prepared in bulk: Beads were treated with Hydrolysis Buffer, washed with 1 ml High Salt Wash Buffer, and washed with 1 ml Binding Buffer as described above in the section *Preparation of streptavidin-coated magnetic beads for P*_*RA1*_ *barcoding assays*. After removing Binding Buffer, the beads were washed twice with 500 μl of Wash Buffer T by transferring to a new tube, washing with rotation for 5 minutes at room temperature, and removing the supernatant. After washing the second time with Wash Buffer T, the beads were resuspended to a concentration of 1 μg/μl in Wash Buffer T, split into 25 or 50 μl aliquots, and stored on ice until use.

### Purification of TECs by selective photo-elution from streptavidin beads

Several protocols for purifying TECs were performed, both for the purpose of protocol development and for assessing the properties of TECs that were prepared in different ways. For simplicity, the final validated protocol is detailed first, and other variations that were performed are then described below with reference to the figure(s) in which each procedure was used. Preparations using the final protocol are shown in Figures 4B, 4D, 5B, S2, S3A, and S3B. All sample handling was performed under low intensity 592 nm amber light until the 365 nm UV irradiation step.

For TEC purification ‘Target DNA’ refers to DNA Template 4 (Table S2), which contains an internal biotin-TEG modification as the RNAP stall site and a P_RA1_ promoter with 5’ PC biotin and C3 Spacer modifications at the 5’ end and dU modifications at positions -13 and -30 of the promoter. ‘Competitor DNA’ refers to DNA Template 5 (Table S2), which contains an internal etheno-dA stall site. For reach transcription reaction, 5 μl of 10 mg/ml Dynabeads™ MyOne™ Streptavidin C1 beads were prepared in advance as described above in the procedure *Preparation of streptavidin beads for TEC purification* and stored on ice at a concentration of 1 μg/μl in 50 μl of 1X Wash Buffer T until use. 25 μl *in vitro* transcription reactions containing 1X Transcription Buffer, 500 μM NTPs, 0.1 mg/ml BSA, 10 nM Target DNA, and 0.016 U/μl *E. coli* RNAP holoenzyme were prepared in a 1.7 ml microcentrifuge tube on ice; at this point the total reaction volume was 20 μl due to the omission of Competitor DNA and 10X Start Solution. Transcription reactions were placed in a dry bath set to 37 °C for 20 minutes to form open promoter complexes. After ∼17 minutes, a 1.7 ml microcentrifuge tube containing 2.5 μl of 150 nM Competitor DNA was placed in the 37 °C dry bath to pre-warm. After open complexes had formed on Target DNA for 20 minutes, the 20 μl transcription reaction was transferred to the tube containing 2.5 μl of 150 nM of Competitor DNA; the final concentration of Competitor DNA in the full 25 μl reaction volume was 15 nM. The transcription reaction was returned to 37 °C for an additional 20 minutes so that free RNAP holoenzyme formed open complexes with the Competitor DNA. Dilution Buffer (1X Transcription Buffer supplemented with 10 μg/ml rifampicin and 0.1% Triton X-100) and streptavidin beads were removed from ice and kept at room temperature at this time. After ∼17 minutes, the streptavidin beads were pipetted to resuspend settled beads. After open complexes had formed on competitor DNA for 20 minutes, single-round transcription was initiated by adding 2.5 μl of freshly prepared 10X Start Solution, incubated at 37 °C for 2 minutes, and gently diluted into 425 μl of room temperature Dilution Buffer by pipetting. Dilution Buffer contains rifampicin to maintain single-round transcription conditions during bead binding. The purpose of diluting the reaction was to minimize the occurrence of bead cross-linking by DNA templates in which both biotin modifications were exposed; when bead binding was performed using a sample volume of 25 μl, substantial bead clumping was observed. No observable clumping occurred when the binding reaction was diluted 10- or 20-fold. The diluted 450 μl transcription reaction was then gently mixed with 50 μl of 1 μg/μl streptavidin beads by pipetting. The bead binding reaction was incubated in the dark at room temperature with rotation for 1 hour. After 1 hour, the bead binding mixture was spun briefly in a Labnet Prism mini centrifuge (Labnet International) by flicking the switch on and off so that liquid was removed from the tube cap. The 1.7 ml tube containing the bead binding reaction was placed on a magnet stand for at least 2 minutes to pellet the streptavidin beads on the tube wall, and the supernatant was carefully removed; this supernatant, which contains any reaction components that did not bind the beads including virtually all Competitor DNA, is referred to as fraction S1. The 1.7 ml tube containing the beads was removed from the magnet stand, the beads were gently resuspended in 500 μl of Wash Buffer TM (1X Transcription Buffer supplemented with 1 mM MgCl_2_ and 0.1% Triton X-100) by pipetting, and the sample was returned to the magnet stand for two minutes to pellet the streptavidin beads before removing the supernatant; this supernatant, which contains residual reactions components, is referred to as fraction W. The streptavidin beads were then gently resuspended in 25 μl of Wash Buffer TM by pipetting so that the bead concentration was 2 μg/μl, and placed in a custom-built 365 nm LED irradiator for 1.7 ml microcentrifuge tubes (Fig. S1, see *Assembly and validation of a 365 nm microcentrifuge tube irradiator* below for details) and exposed to ∼10 mW/cm^2^ 365 nm UV light from four directions for 5 minutes. After irradiation, the bead mixture was returned to the magnet stand for 1 minute before collecting the supernatant; the pelleted beads, which contain DNA without a TEC and any TECs that were not eluted by 365 nm UV irradiation, are referred to as fraction P. The collected supernatant, which contains purified TECs, is referred to as fraction S2.

Several variations on this protocol were performed. In the proof-of-principle experiment in Figure 3D, the transcription reaction included 5 nM Target DNA and 0.024 U/μl RNAP holoenzyme, Competitor DNA was not included, half the amount of streptavidin beads (25 μg vs. 50 μg) were used, and the volumes used for the bead binding and wash steps were halved (250 μl vs. 500 μl). In Figures 4B, 5B, and S2, TECs were prepared with variable RNAP, Target DNA, and Competitor DNA concentrations. The conditions used for each preparation are indicated by the gel annotations. All samples that used 5 nM template were prepared with the protocol modifications described for the TEC preparation in Figure 3D.

### Analysis of TEC purification fractions by denaturing PAGE

TEC purification fractions were prepared for denaturing PAGE as follows: Fractions S1 and W were mixed with 5 μl of 0.5 M EDTA (pH 8.0). To recover immobilized DNA in Fraction P, the bead pellet was resuspended in 25 μl of 95% formamide and 10 mM EDTA, heated at 100 °C for five minutes, placed on a magnet stand, and the supernatant was collected and mixed with 125 μl of Stop Solution. Fraction S2 was mixed with 125 μl of Stop Solution. The volumes of Fractions P and S2 were then raised to match the volume of Fractions S1 and W (250 μl in Fig. 3D or 500 μl in Fig. 4D) by adding 1X Transcription Buffer. The fractions were extracted by adding an equal volume of Tris (pH 8) buffered phenol:chloroform:isoamyl alcohol (25:24:1, v/v), mixing by vortexing and inversion, and centrifuging at 18,500 x g and 4 °C for five minutes. The aqueous phase was collected and transferred to a new tube. When the volume of the extracted samples was 250 μl, nucleic acids were precipitated by adding 25 μl 3 M NaOAc (pH 5.5), 750 μl 100% ethanol, and 1.5 μl GlycoBlue™ Coprecipitant to each sample and chilling at -70 °C for 30 minutes. When the volume of the extracted samples was 500 μl, nucleic acids were precipitated by adding 50 μl 3 M NaOAc (pH 5.5), 350 μl 100% isopropanol, and 1.5 μl GlycoBlue™ Coprecipitant to each sample and chilling on ice for 30 minutes. The samples were centrifuged at 18,500 x g and 4 °C for 30 minutes and the supernatant was removed. When isopropanol precipitations were performed, the pellet was washed once by adding 1 ml of cold 70% (v/v) ethanol, inverting the tube several times, centrifuging at 18,500 x g and 4 °C for 2 minutes, and removing the supernatant. After removing residual ethanol, the pellet was dissolved in 16 μl of BPB Only Formamide Loading Dye and the entire sample was loaded on the gel. Urea-Page was performed as described above in the section *Denaturing Urea-PAGE*.

### Analysis of purified TECs by EMSA

After purifying TECs as described above, 15 μl of fraction S2, which contains purified TECs, was gently mixed with 3 μl of 6X BPB Only Native DNA Loading Dye. Samples were loaded on a 0.5X TBE 5% polyacrylamide gel prepared using ProtoGel acrylamide. Gels were immersed up to the wells in 0.5X TBE to minimize heating and run at room temperature at 45V for approximately 1.5 hours before being stained using 1X SYBR Gold in 0.5X TBE and scanned on a Sapphire Biomolecular imager using the 488 nM/518BP22 setting.

### USER^®^ protection assay

For the USER^®^ protection assay, TECs were prepared in parallel using the final validated protocol (10 nM Target DNA (DNA Template 4 in Table S2), 0.016 U/μl RNAP holoenzyme, and 15 nM Competitor DNA (DNA Template 5 in Table S2) and conditions that mostly yielded a slow-migrating product when assessed by EMSA (5 nM Target DNA, 0.024 U/μl RNAP holoenzyme, and no Competitor DNA). One modification was made to the TEC purification protocol: Wash Buffer TM was supplemented with 10 μg/ml rifampicin so that the TEC preparations contained 1X Transcription Buffer, 1 mM MgCl_2_, and 10 μg/ml rifampicin; this prevents activity by any remaining open complexes during the USER^®^ digestion, which requires ATP and magnesium. During the bead binding step of the TEC purification protocol, several control reactions were prepared. All 25 μl control reactions contained 1X Transcription Buffer, 0.1 mg/ml BSA, and 5 nM DNA Template 4 (Table S2); the +RNAP control reaction contained 0.024 U/μl RNAP holoenzyme, which saturates the P_RA1_ promoter with open complexes (Fig. 3A). At the time of preparation, control reactions were 20 μl due to the omission of 10X rifampicin and 10X T4 DNA Ligase Buffer.

Upon elution from streptavidin beads, 25 μl aliquots of purified TECs were mixed with 2.5 μl of 10X T4 DNA Ligase Buffer and either placed directly in a dry bath set to 37 °C (-USER^®^ samples) or mixed with 0.5 μl of Thermolabile USER^®^ II Enzyme before being placed at 37 °C (+USER^®^ samples). In both cases, purified TECs were incubated at 37 °C for 30 minutes, which far exceeds the time required for complete USER^®^ digestion (<5 minutes). Control reactions were incubated at 37 °C for 15 minutes to form open complexes. Rifampicin was added to 10 μg/ml and the reactions were incubated at 37 °C for 5 additional minutes before the addition of 2.5 μl of 10X T4 DNA Ligase Buffer and, in the case of the +USER^®^ samples, 0.5 μl of Thermolabile USER^®^ II Enzyme. The control reactions were incubated 37 °C for 30 minutes. USER^®^ digestion was stopped by the addition of 125 μl of Stop Solution, and the samples were processed for denaturing PAGE by phenol:chloroform extraction and ethanol precipitation as described in the section *P*_*RA1*_ *barcoding validation assays*, resuspended in 16 μl of BPB Only Formamide Loading Dye and the entire sample was used for denaturing PAGE.

### TEC degradation assay

TECs were purified as described above; 1 reaction volume was removed from the *in vitro* transcription master mix before the addition of *E. coli* RNAP holoenzyme and kept on ice as a DNA only control. Purified TECs were split into 25 μl aliquots and kept at room temperature. When included in the reaction, 1 μl of Thermolabile Proteinase K was added and mixed with the sample immediately before the sample was placed at 37 °C. All samples were incubated at 37 °C for 30 minutes, and one sample was heat inactivated by incubating at 65 °C in a thermal cycler for 10 minutes. The time course was structured so that all sample handling reached completion simultaneously. 15 μl of each sample was gently mixed with 3 μl 6X BPB Only Native DNA Loading Dye and assessed by EMSA as described above in the section *Analysis of purified TECs by EMSA*.

### P_RA1_ barcoding of DNA from purified TECs

TECs were prepared as described above, except that 50 μl transcription reactions were used, and the volume of every step was adjusted accordingly. Purified TECs were pooled. A 15 μl aliquot was removed and assessed by EMSA as described above in the section *Analysis of purified TECs by EMSA*, and a 25 μl ‘Input’ fraction was mixed with 125 μl of Stop Solution, phenol:chloform extracted and set up for ethanol precipitation as described above in the section *P*_*RA1*_ *barcoding validation assays*. All subsequent incubations were performed in a thermal cycler. 150 μl of the remaining sample was mixed with 6 μl (1 μl per sample volume) of Thermolabile Proteinase K (NEB), incubated at 37 °C for 30 minutes, and then at 65 °C for 20 minutes. 3 μl (0.5 μl per sample volume) of RNase I_f_ (NEB) was added and the sample was incubated at 37 °C for 15 minutes, and then at 70 °C for 20 minutes. Together these steps degraded RNAP and RNA, and the sample was stored at -20 °C overnight. The next day, 26.5 μl was removed and mixed with 125 μl of Stop Solution. The remaining sample was mixed with 12.5 μl (2.5 μl per sample volume) of 10X T4 DNA ligase Buffer and 2.5 μl (0.5 μl per sample volume) of Thermolablie USER^®^ II Enzyme and processed exactly as described above in the section *P*_*RA1*_ *barcoding validation assays* through the Klenow Fragment primer extension step. After each enzymatic processing step a reaction fraction (with volume adjusted for added components) was mixed with 125 μl of Stop Solution. All intermediate fractions were phenol:chloroform extracted and ethanol precipitated as described above in the section *P*_*RA1*_ *barcoding validation assays*, and the pellet was resuspended in 16 μl of BPB Only Formamide Loading Dye for denaturing PAGE. After all intermediate fractions were collected, this procedure yielded two reaction volumes of bead-immobilized barcoded DNA. The beads were washed once with 1 ml of 10 mM Tris-HCl (pH 8.0) supplemented with 0.05% Triton X-100, and resuspended in 25 μl of this same buffer for storage at -20 °C.

PCR of the barcoded DNA was performed in 25 μl reactions in 200 μl thin-walled PCR tubes containing 1X Q5^®^ Buffer, 1X Q5^®^ GC Enhancer, 0.2 mM dNTPs, 0.25 μM primer RPI1 (Table S1), 0.25 μM primer dRP1_NoMod.R (Table S1), 2 μl bead-immobilized barcoded DNA, and 0.02 U/μl Q5^®^ High-Fidelity DNA Polymerase (NEB). Reactions were amplified using the thermal cycling program: 98 °C for 30 s, [98 °C for 10 s, 69 °C for 20 s, 72 °C for 20 s] x N cycles (where N = 3, 4, or 5 cycles as indicated in Fig. S3B), hold at 12 °C. After amplification, reactions were transferred to 1.7 ml microcentrifuge tubes and placed on a magnet stand to pellet the beads. The supernatant was collected, and 5 μl of supernatant was mixed with 1 μl of 6X BPB Only SDS DNA Loading Dye (30% (v/v) glycerol, 10 mM Tris-HCl (pH 8.0), 0.48% (w/v) SDS, 0.01% (w/v) bromophenol blue) for electrophoresis as described above in the section *Native PAGE for the analysis of purified DNA*.

### Assembly and validation of a 365 nm microcentrifuge tube irradiator

To enable efficient 365 nm UV-induced cleavage of 5’ PC biotin in a magnetic bead mixture (which can reduce photocleavage efficiency due to light scattering), a custom 1.7 ml microcentrifuge tube irradiator was assembled using 365 nm realUV™ LED Light Strips (Waveform Lighting, Product # 7021.65). The irradiator was designed such that each tube was irradiated by four individual segments of the LED light strip (3 LEDs each, ∼10 mW/cm^2^ from ∼1 cm away from the tube) simultaneously. The irradiator was constructed using the 365 nm LED strips above, LED Strip to Strip Solderless Connectors (Waveform Lighting, Product # 3071), a Female DC Barrel Jack Plug Adapter (Waveform Lighting, Product # 7094), the tube-holder insert from a Beta Box for 1.5 ml microcentrifuge tubes (Fisher Scientific, Cat # 12-009-14), a Desktop AC Adapter (Mean Well, Cat # GST60A12-P1J), and a NEMA 5-15P to IEC320C13 Universal Power Cord (C2G, Product #03129). 3M 12mm VHB Double Sided Foam Adhesive Tape 5952 and hardware including corner brackets, machine screws, washers, and T plates were used to mount the LED strips to the tube-holder insert. A UVA/B light meter calibrated to 365 nm (General, Item # UV513AB) was used to assess 365 nm UV intensity.

To assess 365 nm UV-induced cleavage of 5’ PC biotin, 25 μl reactions containing 1X Transcription Buffer, 0.1 mg/ml BSA, and 5 nM primer PRA1_2dU_PCbio.F (Table S1) were mixed with streptavidin coated magnetic beads (prepared as described above in *Preparation of streptavidin beads for TEC Purification*) to 2 μg/μl or 1 μg/μl and incubated at room temperature with rotation for 30 minutes. The supernatant from this binding reaction, which contains any oligonucleotide that did not bind the beads, was discarded and the beads were resuspended in 25 μl of Wash Buffer T and exposed to 365 nm UV light at 10 mW/cm^2^ from four directions for five minutes. The samples were placed on a magnet stand and the supernatant was collected in 125 μl of Stop Solution. To recover immobilized DNA, the bead pellet was resuspended in 25 μl of 95% formamide and 10 mM EDTA, heated at 100 °C for five minutes, placed on a magnet stand, and the supernatant was collected and added to 125 μl of Stop Solution. Reactions were processed for denaturing Urea-PAGE by phenol:chloroform extraction and ethanol precipitation as described above in the section *P*_*RA1*_ *barcoding validation assays* and resuspended in 7 μl BPB Only Formamide Loading Dye. Urea-Page was performed as described above in the section *Denaturing Urea-PAGE*.

### Quantification

Quantification of band intensity was performed using ImageJ 1.51s by plotting each lane, drawing a line at the base of each peak to subtract background, and determining the area of the closed peak. Because EMSA bands had long trails at the edge of the lane that could overlap for multiple bands, TEC purity was calculated using the internal width of the lane that did not include the trails at the edge.

### Reproducibility of the methods

Key procedures for TEC purification and P_RA1_ barcoding were performed at least five times during protocol development and as controls for validation assays, some of which are presented here. Of particular importance, EMSAs of four independent TEC preparations that were purified using the final validated protocol (detailed below under *Purification of TECs by selective photo-elution from streptavidin beads*) can be found in Figures 4B, S2, S3A, and S3B, and stepwise verification of the P_RA1_ barcoding procedure is shown in Figures 2C, 2D, 2E, and S3B.

## Data availability

All data are contained in the manuscript as plotted values or representative gels. Source files in.tif format are available from the corresponding author (E.J.S.) upon request.

## Supporting information

This article contains supporting information.

## Acknowledgements

I am grateful to J.B. Lucks (Northwestern University) and J.T. Lis (Cornell University) for allowing me to conduct preliminary experiments for this work in their laboratories.

## Author Contributions

E.J.S. is responsible for all aspects of this work.

## Funding

This work was supported by startup funding from the University at Buffalo (to E.J.S.), an Arnold O. Beckman postdoctoral fellowship (to E.J.S.), and a Cornell University Genomics Innovation Hub Seed Grant (to J.T.L. and E.J.S.).

## Conflict of Interest

The author declares that he has no conflicts of interest with the contents of this article.

## Abbreviations

The abbreviations used are:

RNAP: RNA polymerase
TEC: transcription elongation complex
TEG: triethylene glycol
PC: photocleavable
dU: deoxyuridine
USER: uracil specific excision reagent
etheno-dA: 1,N^6^-etheno-2’-deoxyadenosine
P: pellet
S: supernatant

---

**Figure S1.**
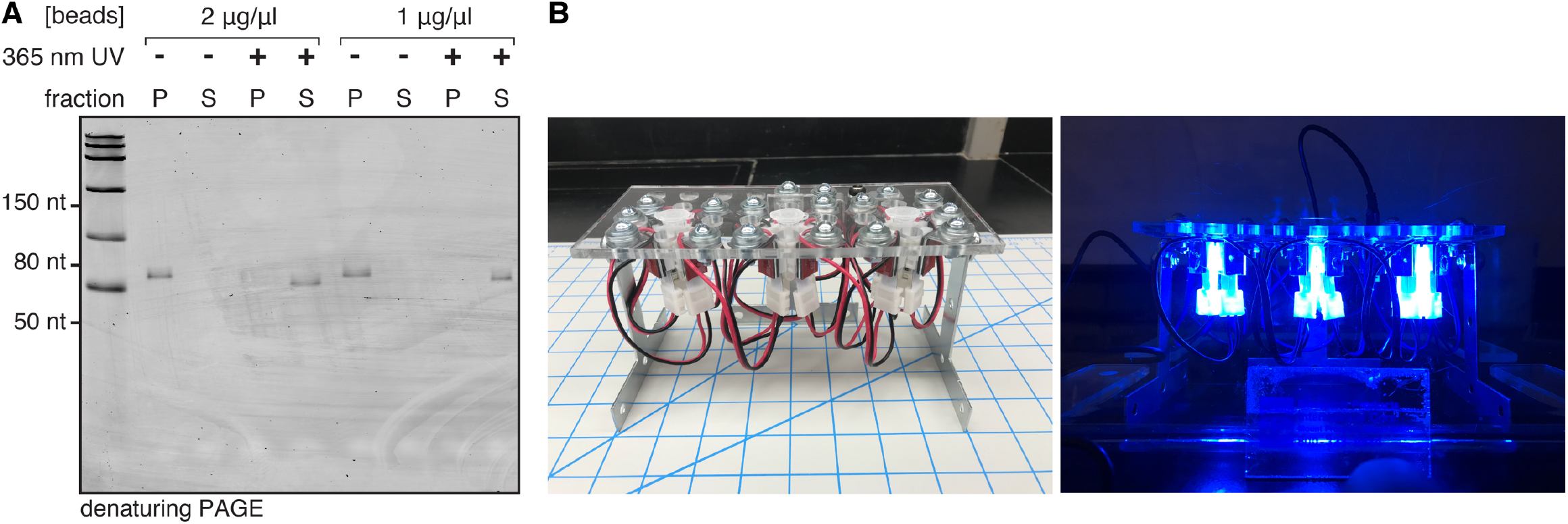
Verification of 5’ PC Biotin photocleavage using a custom 365 nm microcentrifuge tube irradiator. **(A)** Denaturing gel showing efficient 365 nm UV-dependent release of a 5’ PC biotin-modified oligonucleotide from two concentrations of streptavidin-coated magnetic beads. **(B)** Custom built 365 nm tube irradiator.

**Figure S2.**
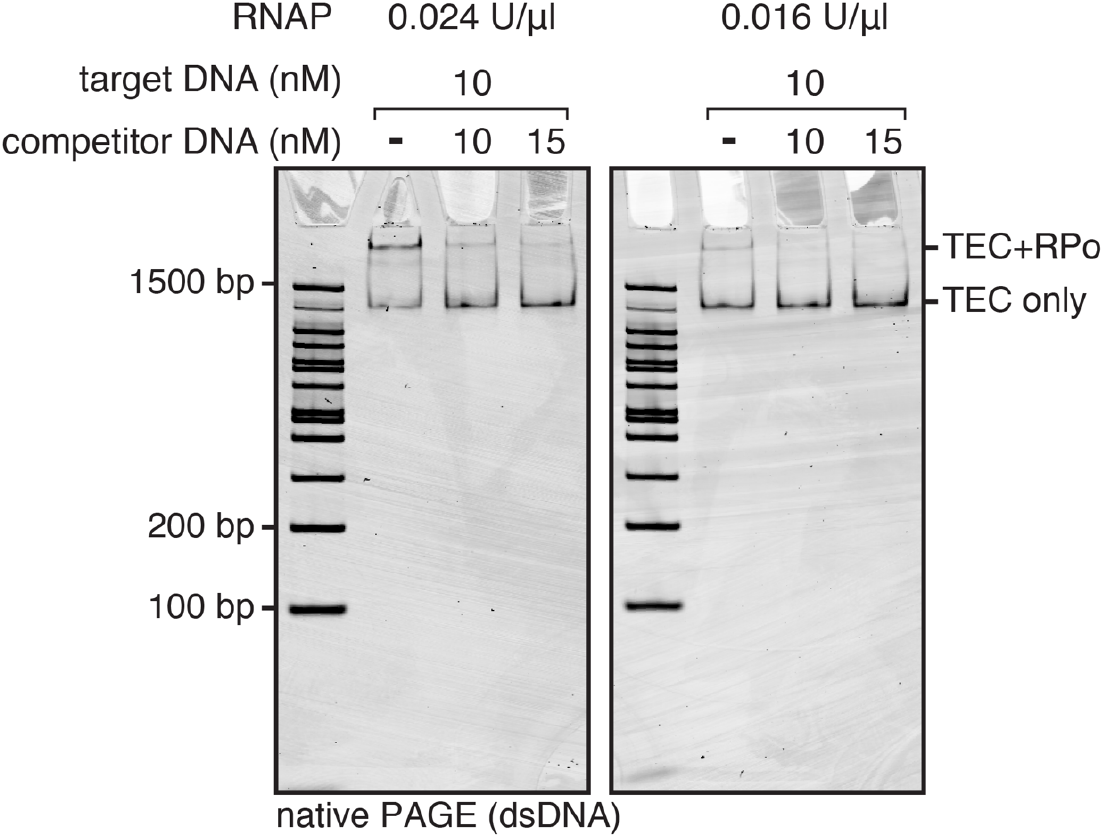
Comparison of TEC purification conditions. EMSA of TECs purified using variable RNAP and competitor DNA concentrations. The gel containing 0.016 U/μl RNAP samples is a replicate of the corresponding gel shown in Figure 4B shown here for comparison to the 0.024 U/μl RNAP samples, and to illustrate the reproducibility of the purification.

**Figure S3.**
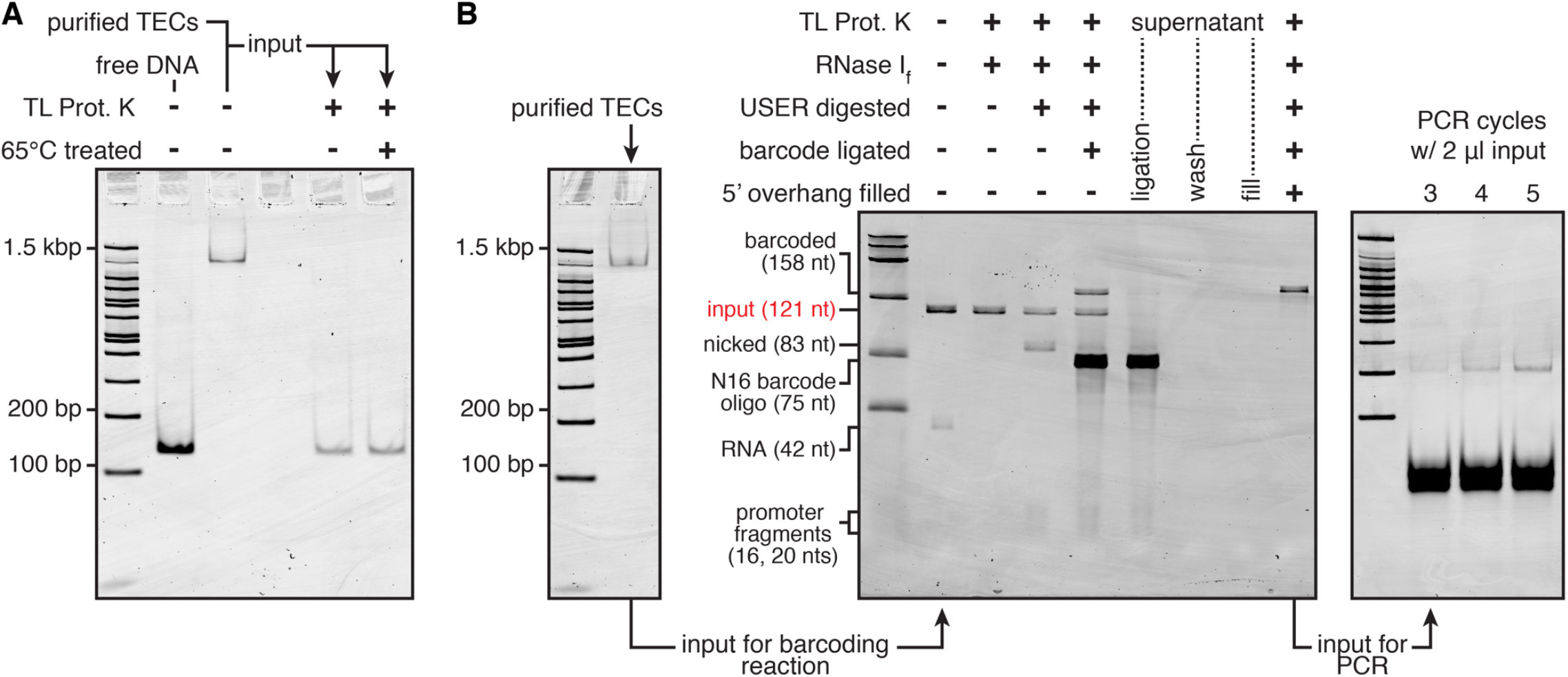
Integration of TEC purification with USER barcoding. **(A)** Degradation of purified TECs using Thermolabile Proteinase K (TL Prot. K) followed by heat treatment at 65 °C. **(B)** Stepwise visualization of the USER barcoding procedure beginning from a single pool of purified TECs.

**Figure S4.**
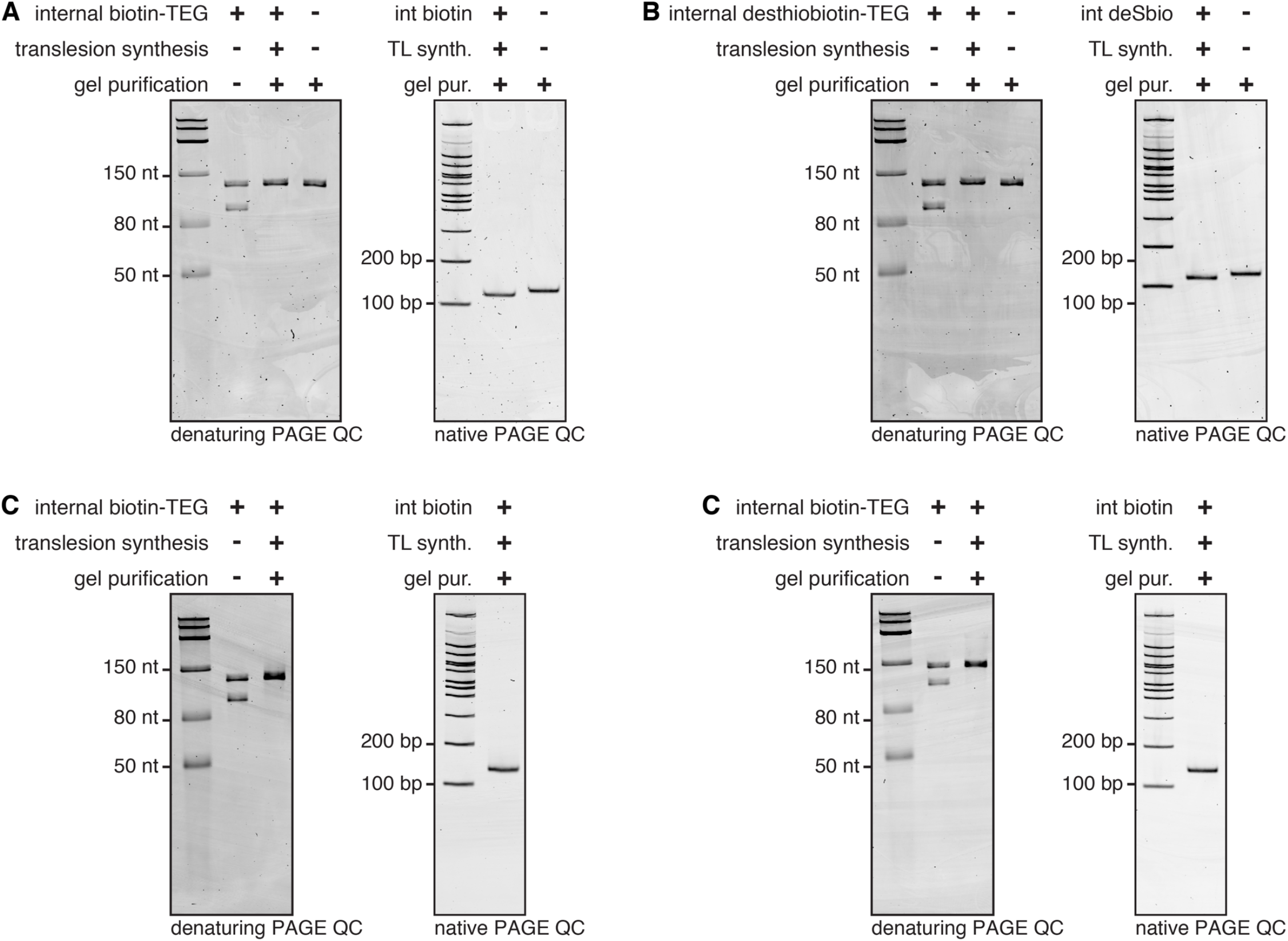
Quality control for internally modified DNA template preparations. Denaturing and non-denaturing quality control of **(A)** DNA template 2, **(B)** DNA template 3, **(C)** DNA template 4, and **(D)** DNA template 5. Details of each DNA template preparation are available in Table S2. In Panels A and B, DNA Template 1 is used as a positive control DNA that did not contain an internal modification in both the denaturing and non-denaturing quality control gels. The presence of an internal modification causes a slight mobility shift relative to DNA without an internal modification.

**Table S1.**
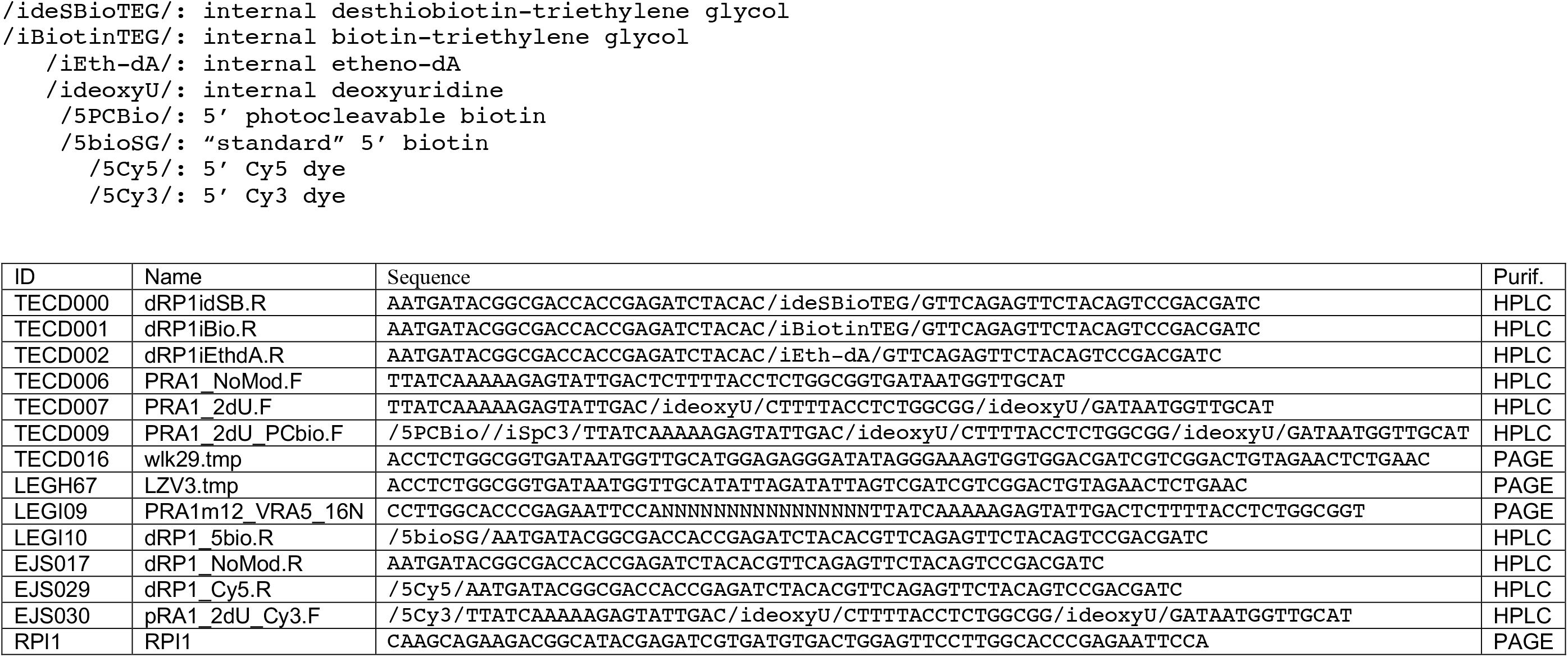
Oligonucleotides used in this study. Below is a table of oligonucleotides used for the preparation of *in vitro* transcription DNA templates. The modification codes defined below are used for compatibility with Integrated DNA Technologies ordering. DNA containing internal biotin-TEG and internal etheno-dA requires an off-catalog order.

**Table S2.** DNA templates prepared for this study. Below is a table of DNA templates that were prepared for this study, including the primers and template oligos used, DNA modifications, the PCR polymerase used, whether translesion synthesis was performed, which reaction clean up protocol was used (see Experimental Procedures), and the figures in which each DNA template was used.

| ID | Fwd Primer | Rev Primer | Template Oligo | Modifications | PCR Polymerase | Translesion Synthesis | Clean Up | Used in Fig(s) |
| --- | --- | --- | --- | --- | --- | --- | --- | --- |
| 1 | TECD007 | LEGI10 | LEGH67 | -13,-30 dU;<br>5' biotin-TEG | Q5U | N/A | Gel Extracted | 2C |
| 2 | TECD007 | TECD001 | LEGH67 | -13,-30 dU;<br>Int biotin-TEG | Q5U | Yes | Gel Extracted | 2D, 2E, 3A, 3B, 4A, S4 |
| 3 | TECD007 | TECD000 | LEGH67 | -13,-30 dU;<br>Int desthiobiotin-TEG | Q5U | Yes | Gel Extracted | 2F, S4 |
| 4 | TECD009 | TECD001 | LEGH67 | 5' PC biotin;<br>Int C3 spacer;<br>-13,-30 dU;<br>Int biotin-TEG | Q5U | Yes | Thermolabile Exol<br>+ PCR clean-up | 3D, 4B, 4D, 5B, S2, S3A, S3B, S4 |
| 5 | TECD006 | TECD002 | TECD016 | Int etheno-dA | Q5 | Yes | Thermolabile Exol<br>+ PCR clean-up | 4B, 4D, 5B, S2, S3A, S3B, S4 |
| 6 | EJS030 | EJS017 | LEGH67 | 5' Cy3<br>-13,-30 dU; | Q5U | N/A | Thermolabile Exol<br>+ PCR clean-up | 4C |
| 7 | TECD007 | EJS029 | TECD016 | -13,-30 dU;<br>5' Cy5 | Q5U | N/A | Thermolabile Exol<br>+ PCR clean-up | 4C |

## Notes

### Competing Interest Statement

The authors have declared no competing interest.

